# ERK3/MAPK6 dictates Cdc42/Rac1 activity and ARP2/3-dependent actin polymerization

**DOI:** 10.1101/2022.10.12.511969

**Authors:** Katarzyna Bogucka-Janczi, Gregory Harms, Mary May-Coissieux, Mohamad Bentires-Alj, Bernd Thiede, Krishnaraj Rajalingam

## Abstract

The actin cytoskeleton is tightly controlled by RhoGTPases, actin binding proteins and nucleation-promoting factors to perform fundamental cellular functions. Here, we show that ERK3, an atypical MAPK, directly acts as a guanine nucleotide exchange factor for Cdc42 and phosphorylates the ARP3 subunit of the ARP2/3 complex at S418 to promote filopodia formation and actin polymerization, respectively. Consistently, depletion of ERK3 prevented both basal and EGF-dependent Rac1 and Cdc42 activation, maintenance of F-actin content, filopodia formation and epithelial cell migration. Further, ERK3 protein binds directly to the purified ARP2/3 complex and augments polymerization of actin *in vitro*. ERK3 kinase activity is required for the formation of actin-rich protrusions in mammalian cells. These findings unveil a fundamentally unique pathway employed by cells to control actin-dependent cellular functions.

## Introduction

Actin is one of the most abundant and highly conserved proteins with over 95% homology among all isoforms^1^. In cells, it is present in globular/monomeric form (G-actin), which can polymerize into branched and elongated filamentous forms (F-actin) in a dynamic, spatially and temporally controlled process^2–5^. The cytoplasmic actin network constitutes an important part of the cytoskeleton, which not only mechanically supports the plasma membrane and gives the cell its shape, but fulfils a variety of other functions: It regulates velocity and directionality of cell migration, enables intracellular signaling and transport, supports cell division and more^6, 7^. Moreover, by branching and bundling of F-actin filaments the actin cytoskeleton can form unique structures such as lamellipodia and filopodia, which in epithelial cells regulate cell polarization and are crucial for the contact with their environment, while in other cell types these actin rich structures control motility, chemotaxis and haptotaxis^8–12^.

To perform its cellular functions, the actin cytoskeleton must be able to rapidly remodel in response to intrinsic and extrinsic cues^13–15^. The spatiotemporally defined branching and elongation of the polymerized actin filaments is achieved by activation of nucleation factors including the Actin-related protein 2 and 3 (ARP2/3) complex, Formins and Spire^16–23^. Actin nucleators facilitate formation of F-actin architectures, branches and bundles, giving rise to specific actin-rich protrusions, thus generating the pushing force required for membrane protrusion during migration^15, 24, 25^. It is well established that ARP2/3–dependent actin polymerization generates branched F-actin network, while Formins generate unbranched filaments and promote growth of actin filaments at the barbed end^21, 26, 27^. Two models of the filopodia initiation emerged, where ARP2/3 and Formins are thought to be involved in the “convergent elongation” and “tip nucleation” model, respectively. However, the proposed models are not mutually exclusive and filopodia formation could engage both Arp2/3 and Formins^27^. Moreover, both *in vivo* and *in vitro* experimental evidence upholds the essential role for ARP2/3 complex-dependent actin nucelation in the formation of filopodia-like protrusions^25, 28–34^.

ARP2/3 complex is a major and highly conserved actin nucleator composed of seven subunits^35, 36^. The structural and biochemical properties of the ARP2/3 complex have been intensively studied *in vitro* and apart from ATP, its activation relies on binding to pre-formed actin filament and nucleation promoting factors (NPFs)^35–40^. The main regulators of the ARP2/3 complex belong to the Wiskott-Aldrich syndrome protein (WASP) family proteins, including WASP-family verprolin-homologous protein (WAVE)^41–43^. WASP and WAVE proteins serve as a link between the actin cytoskeleton and RhoGTPases via direct and intermediate binding to Cell division control 42 (Cdc42) and Ras-related C3 botulin toxin substrate 1 (Rac1), respectively^17, 18^.

The small Rho GTPases belong to the Ras homologous (Rho) superfamily of GTP-binding proteins^44^. They function as binary molecular switches circulating between active (GTP bound) and inactive (GDP bound) state. The on-off regulation of the Rho GTPases is under the control of guanine exchange factors (GEFs) and GTPase activating proteins (GAPs)^44^. GTP-bound active Cdc42 and Rac1 promote the formation of filopodia and lamellipodia, respectively, by activating the NPFs, WASP and WAVE^17, 18^. Recent studies revealed a role for kinases in directly activating Rho GTPases mainly through phosphorylation, but the identity of the relevant kinase(s) remained elusive^45–49^.

ERK3 (MAPK6) is an atypical member of the mitogen-activated protein kinases (MAPKs). Its physiological roles are tissue-specific and can be both dependent and independent of its catalytic activity: Kinase-dependent roles of ERK3 include promoting lung cancer cell migration and invasion, while its function in the regulation of breast cancer cell morphology and migration is shown to be partially kinase-independent^50–53^. Moreover, ERK3 is a labile protein whose rapid turnover has been implicated in cellular differentiation^51, 54^. Although the specific signaling mechanisms involved in the regulation of ERK3 have been studied by us and others and provided tissue-specific insights into this atypical MAPK function^51, 53, 55–58^, many aspects of ERK3 biology such as its role in suppressing melanoma cell growth and invasiveness remain elusive^59^. It is, however, known that ERK3 possesses a single phosphorylation site at serine 189 (S189) within S-E-G activation motif, located in its kinase domain, which was shown to be phosphorylated by group I p-21-activated kinases (PAKs), the direct effectors of Rac1 and Cdc42 GTPases^60–62^.

ERK3 signaling is required for the motility and migration of different cancer cells, yet its direct role in the regulation of the polarized phenotype of the cells and actin cytoskeleton is lacking. Here, we unveil a multi-layered role for ERK3 in regulating actin filament assembly. We demonstrate that ERK3 controls bundling of actin filaments into filopodia via activation of Cdc42. Mechanistically, ERK3 directly binds to Rac1 and Cdc42 in a nucleotide-independent manner and is required for the activity of both RhoGTPases *in vivo.* Furthermore, ERK3 could function as a direct GEF for Cdc42 *in vitro*. In addition, ERK3 stimulates Arp2/3-dependent actin polymerization and maintains F-actin abundance in cells via direct binding and phosphorylation of the ARP3 subunit at S418. While the kinase activity is not required for Cdc42 activation, Rac1 activity partially depends on ERK3 kinase function. The *in vivo* relevance of these ERK3 functions was corroborated by analyses of the motility of tumour cells in orthotopic mammary tumors grown in mice. Together, our results establish a hitherto unknown regulatory pathway directly controlled by ERK3 involving the major signaling molecules Cdc42, Rac1 and Arp2/3 in the modulation of the actin cytoskeleton and cell migration.

## Results

### ERK3 is required for motility of mammary epithelial cells

ERK3 has been shown to regulate mammary epithelial cancer cell migration and metastasis^50, 52^. To uncover the molecular underpinnings of the role of ERK3 in cell migration, we investigated the intracellular localization of ERK3 by immunocytochemical analyses using a validated antibody^51, 52^. Endogenous ERK3 co-localized to the F-actin-rich protrusions in human mammary epithelial cells (HMECs) (Figure 1A). Re-organization of actin protrusions at the leading edge of the cells initiate cell migration^6, 63^, therefore we further analyzed the significance of ERK3 in cancer cell motility by loss of function studies. Depletion of ERK3 reduced speed and displacement length (distance between the start point and the end position of the cell along each axis) of MDA-MB231 cells (Figure 1B-1F and Supplementary Movie 1-4). As shown in Figure 1C, depletion of ERK3 strongly reduced the random cell motility, which remained low over time as indicated by the calculated acceleration (Figure 1D) and concomitantly resulted in a shorter overall displacement length (Figure 1E) with shorter migration track length and average speed (Figure 1-figure supplement 1A and 1B). Considering that responses of tumor cells depend on their microenvironment, we tested the *in vivo* motility of these cells. Intravital imaging of the orthotopic mammary tumors confirmed reduced motility of the ERK3 knockdown MDA-MB231-GFP cells (Figure 1G and Supplementary Movies 5 and 6). Together, these data suggest that ERK3 likely controls actin cytoskeleton dynamics thereby influencing cell shape, motility and polarized migration.

**Figure 1.**
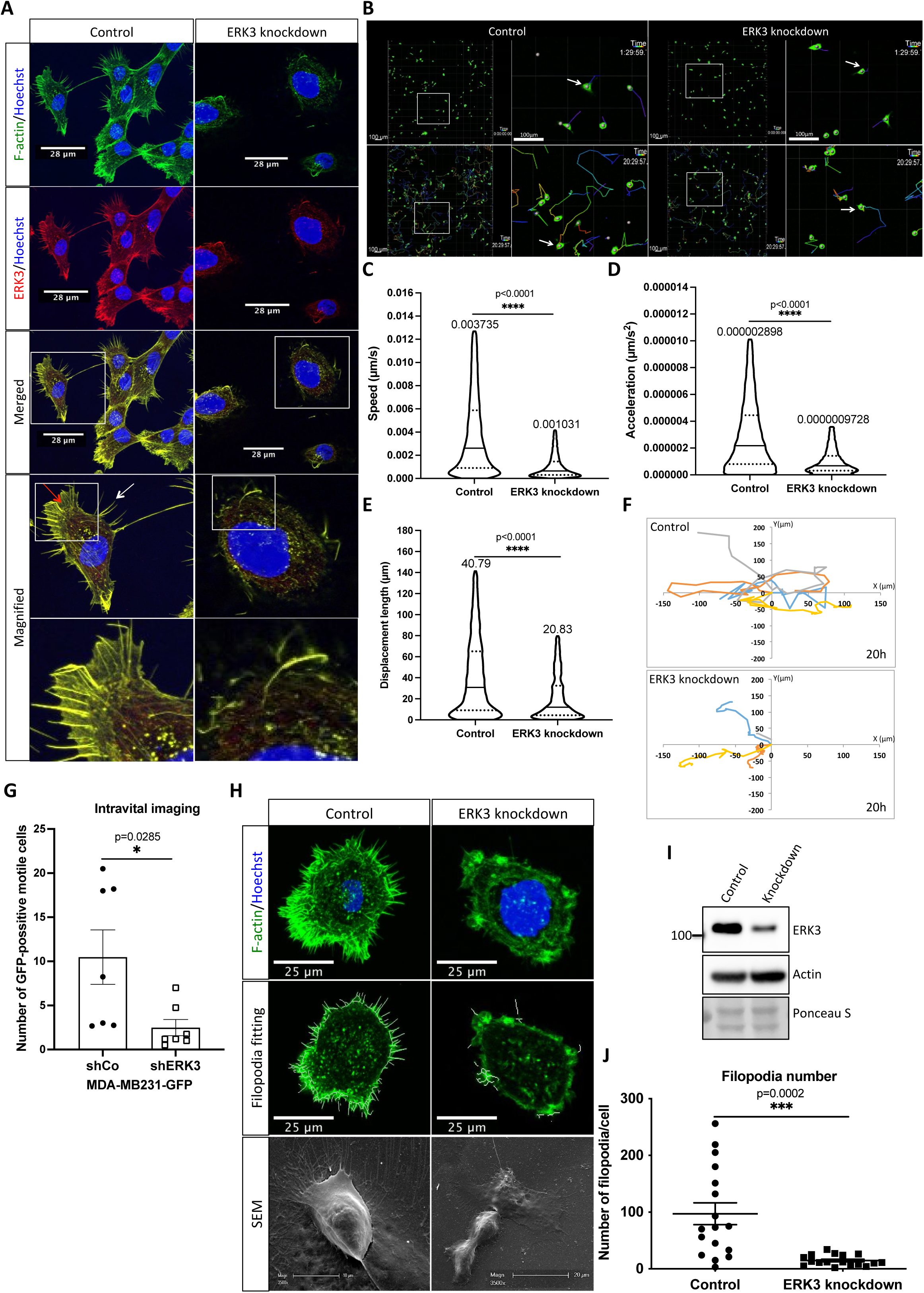
ERK3 localizes to the F-actin-rich protrusions and regulates mammary epithelial cells motility. **(A)** Confocal analyses of control (shCo) and ERK3 knockdown (shERK3) HMECs co-stained with anti-F-actin (green) and anti-ERK3 antibodies (red). Hoechst staining (blue) was used to visualize cell nuclei. Scale bars 28 µm. Higher magnification images of the boxed areas are shown on the right. Red and white arrows exemplify lamellipodia and filopodia, respectively. **(B-F)** Single-cell tracking analyses of control (shCo) and ERK3 knockdown (shERK3) MDA-MB231-GFP cells. **(B)** Representative images of the cell migration assay were taken at the start- (top panel) and end-point (20 h) (lower panel) of the full track length. Random fields were selected, and boxed areas were magnified on the right. Arrows indicate exemplified cell tracking at the beginning and at the end of the tracking. **(C-E)** Violin plots present cell distribution according to the analyzed motility parameters, calculated as described in the methods section. Plotted are median (solid line) and 25%/75% quartiles (dashed lines). **(C)** Speed (µm/s), **(D)** acceleration (µm/s^2^), and **(E)** displacement length (µm) of fourteen thousand six hundred fifty-nine single cells (n=14659) are depicted. Significance was determined using non-parametric Mann-Whitney test. *p<0.0332, **p<0.0021, ***p<0.0002, ****p<0.0001. Analyses of the migration tracks length and average speed are depicted in Figure 1-figure supplement 1A and 1B. **(F)** Cell migration patterns of four randomly selected control and ERK3 knockdown cells were plotted as x-y trajectories over the 20 h tracking. Please see Figure S9 for the Western Blot of the knockdown validation. **(G)** Quantification of the intravital imaging of the control (shCo) and ERK3-knockdown (shERK3) orthotopic mammary tumors (MDA-MB231-GFP). The number of GFP-positive motile cells was quantified as described in the methods section and is presented as mean ± SEM from seven animals (n=7) per condition. Please see representative Supplementary Movie 5 and 6. Chemotactic responses of the ERK3-depleted cells were assessed in the presence of EGF and are presented in Figure 1-figure supplement 1C-1F. ERK3 protein kinetics and stability in response to EGF are depicted in Figure 1-figure supplement 2. **(H-J)** Depletion of ERK3 alters leading edge protrusions in primary mammary epithelial cells**. (H)** Confocal F-actin (green)/Hoechst (blue) and SEM images analyses of control and ERK3 knockdown HMECs are presented after 4 h starvation. For the confocal analyses, exemplified filopodia fitting was included in the middle panel. **(I)** Western Blot analyses presenting ERK3 knockdown efficiency and total levels of β-actin. Ponceau S was used as a loading control. **(J)** Filopodia number was quantified as described in the methods section and is presented as mean ± SEM of 17 single cells (n=17); *p<0.0332, **p<0.0021, ***p<0.0002, ****p<0.0001, unpaired t-test. Effect of S189 phosphorylation of ERK3 on F-actin assembly is presented in Figure 1-figure supplement 3.

### ERK3 is activated and required for EGF-mediated directional migration in epithelial cells

To further elucidate the physiological role of ERK3 in breast epithelial cell motility, we assessed EGF-induced chemotactic responses to evaluate directional migratory properties of control and ERK3-depleted primary HMECs and metastatic MDA-MB231 cells. Interestingly, knockdown of ERK3 significantly attenuated the EGF-induced chemotaxis of both cell types (Figure 1-figure supplement 1C, 1D and 1E, 1F, respectively).

Considering that ERK3 participated in EGF-mediated chemotactic responses of both primary and oncogenic mammary epithelial cells, we tested ERK3 protein kinetics in response to the growth factor. EGF induced serine 189 (S189) phosphorylation of ERK3 at early time points in primary epithelial cells (Figure 1-figure supplement 2A and 2B) and MDA-MB231 cells (Figure 1-figure supplement 2C and 2D). A slight increase in ERK3 protein levels was detected in HMECs upon EGF treatment (Figure 1-figure supplement 2A and 2B). Further cycloheximide (CHX) chase experiments coupled with RT-PCR analyses suggested that EGF treatment did not increase ERK3 protein half-life (Figure 1-figure supplement 2E-2H). Taken together, these data confirmed that ERK3 was activated in response to EGF and required for EGF-mediated directional migration of epithelial cells. We then investigated the significance of S189 phosphorylation of ERK3 on F-actin and formation of actin rich protrusions. ERK3-depleted HMECs cells (shERK3 -3’UTR) presented in Figure 1A were transfected with V5-tagged ERK3 wild type (ERK3 WT) or S189 mutants (ERK3 S189A/ERK3 S189D) and expression of exogenous ERK3 was visualized by V5-tag staining. Co-staining of the cells with phalloidin green did not show any significant differences in F-actin organization upon the overexpression of the S189 phosphorylation mutants as compared to the wild type ERK3 transfected cells (Figure 1-figure supplement 3). Moreover, we observed that exogenous ERK3 localized predominantly in the cytosol as compared to the endogenous ERK3, where we could detect ERK3 localizing at the cell edges and F-actin rich protrusions (Figure 1A). These results prompted us to prefer ERK3 knockdown cells for the study of ERK3-mediated phenotypes.

### ERK3 is required for actin-rich protrusions

In order to migrate, cells acquire morphological asymmetry to drive their locomotion. One of the first steps into breaking spatial symmetry of the cell involves its polarization and formation of membrane extensions^15^. In response to extracellular guidance, cells polarize and extend F-actin-rich protrusions at the leading edge to direct cell migration^63–65^. As shown in Figure 1A, ERK3 co-localized with F-actin at the edge of the cell and at the protruding filopodial spikes. Moreover, knockdown of ERK3 clearly reduced F-actin content and actin-rich protrusions in human mammary primary epithelial cells (Figure 1A). Further analyses of control and ERK3 knockdown HMECs at the single-cell level confirmed that ERK3-depleted cells had limited, diffuse F-actin, concentrated in the cortical areas of the cells, and a significantly decreased number of filopodia (Figure 1G-1J).

### ERK3-dependent regulation of Rac1 and Cdc42 activity

RhoGTPases are the major membrane signaling transmitters that drive polarized cell asymmetry by locally regulating the formation of F-actin-rich protrusions^66, 67^. EGF signaling triggers activation of Rac1 and Cdc42, which link the signal to the actin cytoskeleton leading to the formation of lamellipodia- and filopodia-like protrusions, respectively (Figure 2A)^68^.

**Figure 2.**
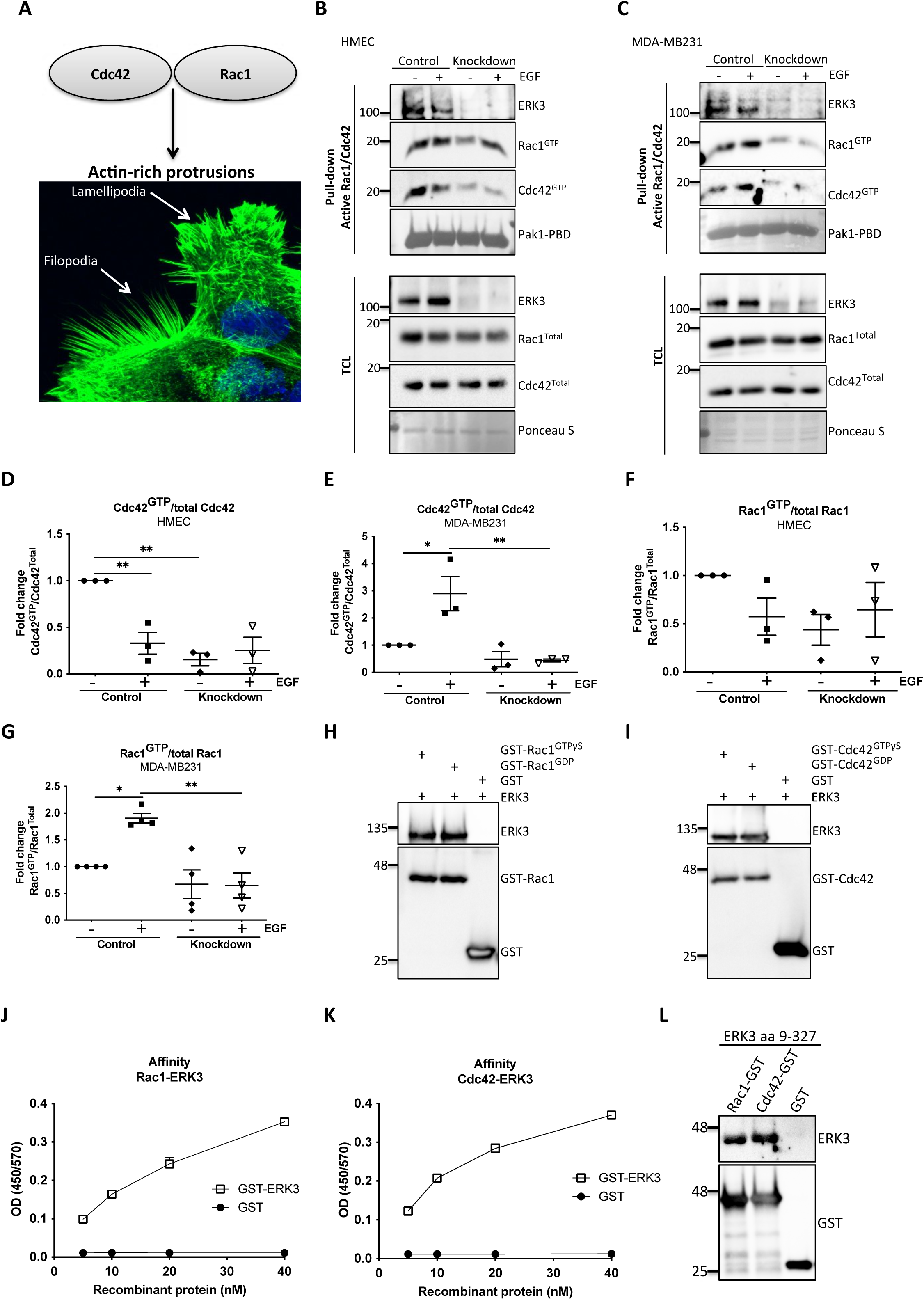
ERK3 binds Rac1 and Cdc42 and regulates their activity. **(A)** Confocal image of phalloidin-stained actin-rich protrusions in HMECs, filopodia and lamellipodia. **(B-G)** *In vivo* pull-down of active Rac1 and Cdc42 from control and ERK3-depleted (knockdown) **(B)** HMECs and **(C)** MDA-MB231 cells in the presence and absence of EGF. Levels of active (GTP-bound) Cdc42 and Rac1 as well as the total protein expression were assessed and quantified. ERK3 knockdown efficiency was verified in the total cell lysate (TCL) as well as the co-immunoprecipitation of ERK3 with active Rac1 and Cdc42. Relative levels of **(D-E)** active Cdc42 and **(F-G)** Rac1 were calculated with respect to the total protein levels and are presented as mean ± SEM from minimum three (n=3) independent experiments; *p<0.0332, **p<0.0021, ***p<0.0002, ****p<0.0001, one-way ANOVA, Tukey’s post-test. ERK3-dependent regulation of the Rac1 and Cdc42 activity was assessed in multiple cell types and data are presented in Figure 2-figure supplement 1 and 2. **(H-I)** *In vitro* interaction between ERK3 and GDP/GTP bound **(H)** Rac1 and **(I)** Cdc42 was assessed, GST was used as a negative control. Pull-down efficiency was assessed with GST antibody and levels of bound ERK3 were verified. The nucleotide-loading status of the purified Rac1 and Cdc42 proteins is presented in Figure 2-figure supplement 3. **(J-K)** Concentration-dependent binding affinity of ERK3-GST protein to the **(J)** Rac1 and **(K)** Cdc42 was determined. Interacting ERK3/GST proteins were used at 5, 10, 20 and 40 nM concentrations. Data from three independent experiments run in triplicates are presented as mean ± SEM. **(L)** ERK3 kinase domain (aa 9-327) binds to Rac1 and Cdc42. Representative analysis of the *in vitro* GST pull-down of Rac1 and Cdc42 and the interaction with the ERK3 kinase domain recombinant protein. Binding affinity of ERK3 (aa 9-327) with Pak1-PBD was verified and is presented in Figure 2-figure supplement 4.

Considering that loss of ERK3 significantly affected F-actin distribution and filopodia formation in primary mammary epithelial cells (Figure 1A and 1G-J), we tested the activity of the two major regulators of actin assembly, Cdc42 and Rac1 (Figure 2 B-G). ERK3 knockdown significantly decreased levels of both basal and EGF-induced GTP-bound Cdc42 and Rac1 in primary (HMEC) (Figure 2B, D, F) and triple negative breast cancer (MDA-MB231) mammary epithelial cells (Figure 2C, E, G), respectively. HMECs were cultured in the growth factor/hormone-enriched medium, including EGF. Therefore, we withdrew the supplements to test the effect of EGF alone on the activity of Rac1 and Cdc42 in these cells and compared the responses with those of the cancer cells. Interestingly, we detected high Rac1 and Cdc42 levels at steady state in HMEC cells, which underwent a rapid activation-inactivation cycle after starvation and restimulation with EGF alone. The inhibitory effect on Rac1 was not as striking as with Cdc42 (Figure 2D-G). Intriguingly, we could observe co-precipitation of ERK3 protein with the active RhoGTPases at endogenous levels suggesting that these proteins exist in a multimeric complex (Figure 2B-2C). We expanded these analyses to different cell types and consistently found that depletion of ERK3 reduces the activity of Rac1 and Cdc42 and that the active RhoGTPases specifically co-precipitated endogenous ERK3 (Figure 2-figure supplement 1 and 2). Interestingly, the effect of ERK3 on Rac1 and Cdc42 activity is either EGF-dependent (MCF7 cells (Figure 2-figure supplement 1A-1C), Calu1 cells (Figure 2-figure supplemnet 2A-2C), –independent (MDA-MB468 cells (Figure 2-figure supplement 1D-1F) or occurs both in resting and EGF-treated cells (HT-29 (Figure 2-figure supplement 2D-2F). Apart from cancer cells, we were also able to confirm that ERK3 sustains the activity of Rac1 and Cdc42 in primary human umbilical vein endothelial cells, HUVECs (Figure 2-figure supplement 2G-2I).

### ERK3 directly binds to Rac1 and Cdc42

In light of these observations, we tested whether any direct interaction existed between these RhoGTPases and ERK3. By employing purified full-length recombinant proteins, we found that ERK3 directly bound to Rac1 and Cdc42 in a nucleotide-independent manner (Figure 2H and 2I). The nucleotide-loading status of the purified Rac1 and Cdc42 proteins was simultaneously assessed by GST-Pak1-PBD pull-down assay followed by immunoblots (Figure 2-figure supplement 3). To further assess the affinity of the binding, we performed concentration-dependent protein binding assays with ELISA as described in the methods section. Binding of full-length ERK3 to Rac1 and Cdc42 could be detected at a low nanomolar range (5 nM) (Figure 2J and 2K). Next, we tested whether the kinase domain of ERK3 (amino acids (aa) 9-327) would be sufficient for this interaction. *In vitro* GST-pull-down assays confirmed that the kinase domain of ERK3 was sufficient for its binding to Rac1 and Cdc42 (Figure 2L, Figure 2-figure supplement 4).

### ERK3 functions as a GEF for Cdc42

The intrinsic GDP-GTP exchange of RhoGTPases is a slow process, which is stimulated *in vivo* by GEFs (Figure 3A)^44^. To examine the ability of the ERK3 kinase domain to stimulate GDP-GTP exchange, we incubated GDP-loaded RhoGTPases with non-hydrolysable GTPγS in the presence or absence of ERK3 kinase domain and quantified the final amount of GTP-bound Cdc42 (Figure 3B and 3C) and Rac1 (Figure 3D and 3E) using GST-Pak1-PBD fusion beads. The ERK3 kinase domain failed to interact directly with the GST-Pak1-PBD protein but bound to active Cdc42 in the same assay confirming that conformationally stable protein is being employed in these assays (Figure 2-figure supplement 4). These experiments showed that although in cultured cells ERK3 regulated the activity of both Cdc42 and Rac1, it only directly stimulated the GDP-GTP exchange of Cdc42 (Figure 3B and 3C). We further corroborated these results by measuring GEF activity of the ERK3 kinase domain *in vitro* by employing a second fluorophore-based assay. The results confirmed those of the previous assay, as addition of ERK3 stimulated GTP-loading on Cdc42 (Figure 3F), but not on Rac1 (Figure 3G). Notably, ERK3 exerted a more potent and long-lasting effect than Dbl’s Big Sister (Dbs), a well-established GEF for Cdc42 and RhoA proteins (Figure 3F)^69^.

**Figure 3.**
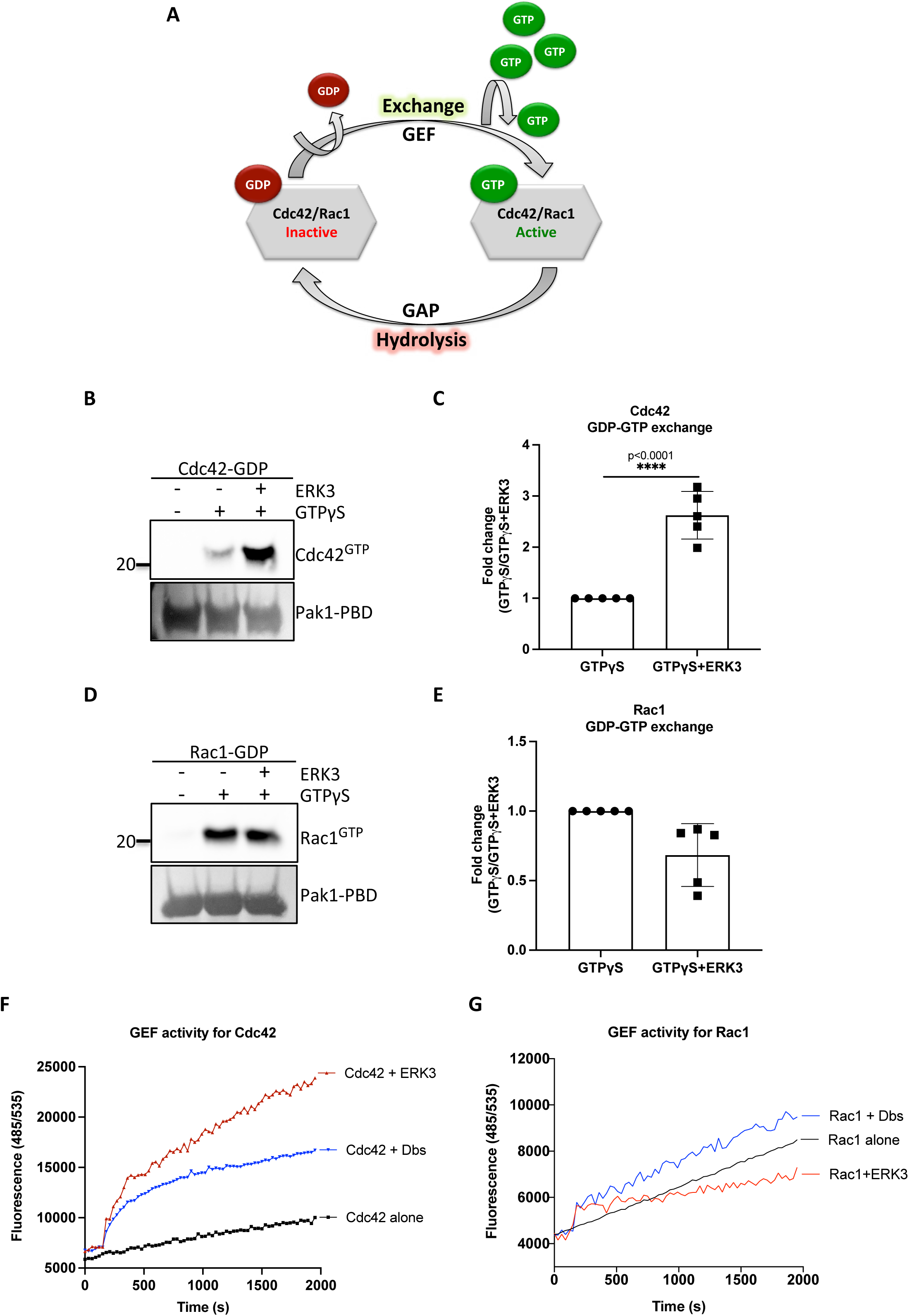
ERK3 kinase domain binds to Rac1 and Cdc42 and facilitates nucleotide exchange on Cdc42. **(A)** Schematic representation of the GDP-GTP nucleotide exchange on Cdc42 and Rac1. Guanine exchange factor (GEF) facilitates the GDP-GTP exchange on the RhoGTPases, resulting in the GTP-bound form of the Cdc42 or Rac1. GTPase activating protein (GAP) stimulates GTP hydrolysis. **(B-E)** Role of ERK3 in the GDP-GTP exchange was assessed in the *in vitro* assays as described in the methods section for **(B-C)** Cdc42 and **(D-E)** Rac1. **(B)** and **(D)** Representative Western Blot analyses of active Rac1 and Cdc42 pull-down, respectively, using Pak1-PBD fusion beads. Levels of active RhoGTPases were detected using Rac1-and Cdc42-specific antibodies. Levels of Pak1-PBD protein were detected by Ponceau S staining and used for the quantification presented in **(C)** and **(E)**. Fold change in GTP-loading was calculated by normalization of the signal obtained for the samples with GTPγS in the presence of ERK3 with samples incubated with GTPγS alone and is presented as mean ± SEM from (D) five (n=5) independent experiments; *p<0.0332, **p<0.0021, ***p<0.0002, ****p<0.0001, unpaired t-test. **(F-G)** *In vitro* RhoGEF activity assay was performed to assess guanine nucleotide exchange activity on **(F)** Cdc42 and **(G)** Rac1 in the presence and absence of recombinant ERK3 protein (9-327 aa). After six initial readings, Dbs-GEF or ERK3 protein were added at 0.5 µM final concentration. GEF activity was expressed as mean relative fluorescence units (RFU) from at least three independent experiments. RhoGEF Dbs protein was used as a positive control. *In vitro* GEF activity of full-length ERK3 vs. ERK3 kinase domain towards Cdc42 was compared as is presented in Figure 3-figure supplement 1.

To assess whether full-length ERK3 could potentiate the effect in terms of activation of Cdc42, we compared GEF activity of the ERK3 kinase domain with that of the full-length protein. Interestingly, full-length ERK3 exhibited weaker GEF activity *in vitro* than the kinase domain of ERK3 (Figure 3-figure supplement 1). The observed *in vitro* discrepancy between the kinase domain and the full-length protein could be attributed to differences in posttranslational modifications due to different production methods of the two recombinant proteins, which might affect conformation and/or activity. The full-length protein employed in these assays was purified from Sf9 cells, while the kinase domain was purified from bacterial cells and therefore lacked – among other things - Ser189 phosphorylation and thus kinase activity *in vitro* (data not shown, also see the methods section). We next examined whether ERK3 is required for the localization and activity of Cdc42 at the plasma membrane (PM). Cellular fractionation assays revealed that knockdown of ERK3 does not disrupt the localization of Cdc42 and Rac1 to the PM (Figure 4A-4C). On the contrary, the PM fraction from ERK3-depleted HMECs had more total Cdc42 and Rac1 than control cells. Interestingly, subsequent pull-down of active Cdc42/Rac1 from the PM fraction revealed that although both RhoGTPAses are present at the plasma membrane in the absence of ERK3, their activity is significantly reduced (Figure 4B and 4C). These results were corroborated by the colocalization of ERK3 with Cdc42 to the protrusions at the cell leading-edge (Figure 4D-4G) with mean Pearson’s correlation coeficient (PCC (r)) of 0.6638 ± 0.02946 and Spearman’s rank correlation coefficent (SRCC (ρ)): 0.7122 ± 0.02586 (Figure 4F and 4G, respectively).

**Figure 4.**
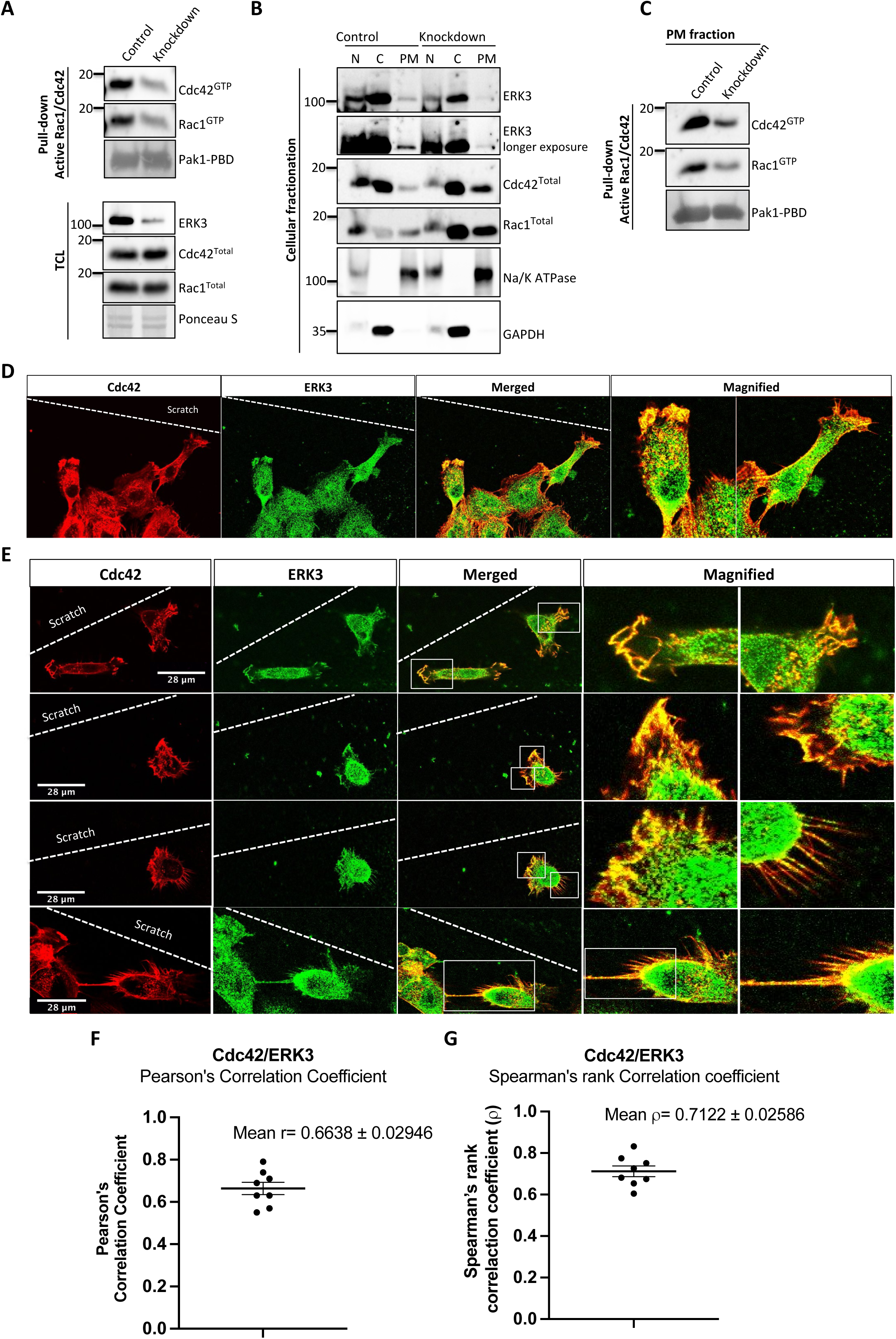
ERK3 colocalizes with Cdc42 in polarized cells and regulates activity of both Cdc42 and Rac1 at the PM. **(A-C)** Cell fractionation for control (siCo) and ERK3-depleted HMECs (siERK3). Fractionation was performed using Minute Plasma Membrane Protein Isolation and Cell Fractionation Kit (Cat# SM-005, Invent Biotechnologies) according to the manufacturer’s instruction. **(A)** Total cell lysate control was used to determine knockdown efficiency and activation status of Rac1 and Cdc42 using active Rac1/Cdc42 pull-down according to manufacturer’s instructions (Cat# 16118/19, ThermoFisher) and as described in the methods section. **(B)** Cellular fractionation was performed. Expression levels of ERK3, Rac1, and Cdc42 were assessed in nuclear (N), cytosolic (C) and plasma membrane (PM) fractions. Na/K ATPases and GAPDH were used as controls for the analysed fractions. **(C)** Active Rac1/Cdc42 pull-down was performed from the isolated PM fraction. **(D-G)** Colocalization of ERK3 and Cdc42 in polarized cells. HMECs were seeded and cultured on cover slips. When cells became around seventy percent confluent, scratch wounds were introduced to the cover slip using a 200 µl tip, medium was exchanged to supplement-free and cells were cultured for additional 6h. Afterwards, cells were fixed and subjected to IF staining as described in the methods with anti-Cdc42 (red) (secondary antibody: Anti-mouse Cy3 (Cat# A10521, ThermoFisher Scientific) and anti-ERK3 (green) (secondary antibody: Alexa Fluor 488 (Cat# A11008, ThermoFisher Scientific) antibodies. Merged images show the colocalization of both proteins in yellow. Magnification of the boxed regions are shown on the right for better visualization. In **(D)** a group of cells at the scratch site is presented and ERK3-Cdc42 colocalization is marked with red arrows at the cell protrusions and with white arrows at the cell body. **(E)** Images representing ERK3-Cdc42 colocalization at a single-cell level at the scratch site. Scale bars 28 µm. **(F-G)** Colocalization of ERK3 and Cdc42 was analyzed as described in the methods and values for **(F)** Pearson’s correlation coefficient as well as the **(G)** Spearman’s rank correlation coefficient are presented for eight randomly selected cells (n=8). Scores above 0 indicate a tendency towards co-localization with a perfect co-localization with a score of 1.

### ERK3 as a nucleation-promoting factor of ARP2/3-dependent actin polymerization

Active Rac1 and Cdc42 induce ARP2/3-dependent actin polymerization by activating their effector NPFs, WASP and WAVE, respectively^17, 18^. Figure 5A depicts a well-described signaling module of ARP2/3-dependent actin nucleation stimulated by Cdc42-activated WASP protein^17, 40, 70–72^.

**Figure 5.**
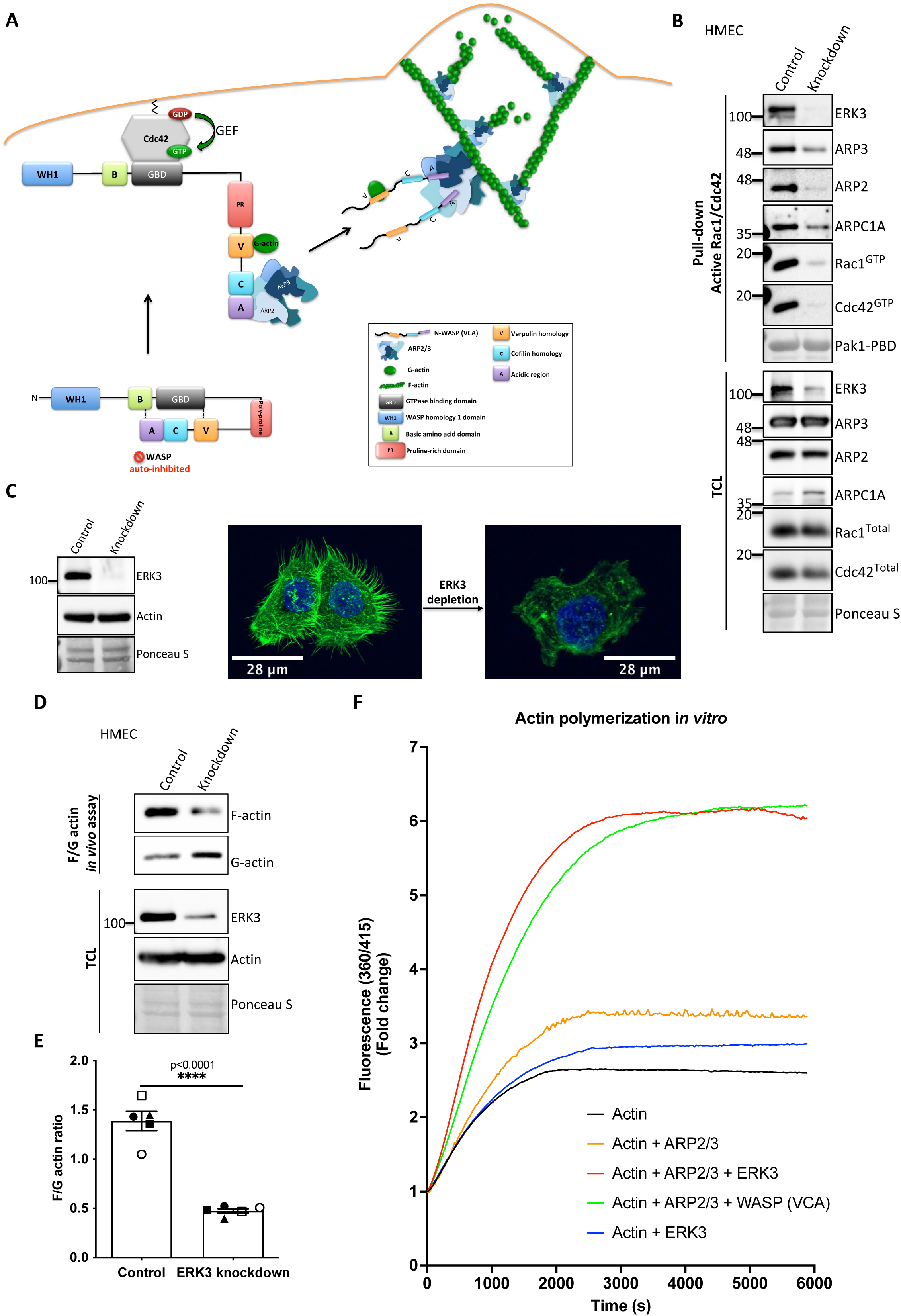
ERK3 acts as a nucleation-promoting factor (NPF) in ARP2/3- dependent actin polymerization *in vitro* and *in vivo*. **(A)** Schematic overview of Cdc42-WASP stimulated ARP2/3-dependent actin polymerization based on the cited literature. The process involves: ARP2/3 complex, WASP (VCA) as nucleation promoting factor, filamentous actin (F-actin) and monomeric actin (G-actin). In the initial step Cdc42 is activated by GEF-catalyzed exchange of GDP to GTP. Active Cdc42 (Cdc42-GTP) binds to the GTP-binding domain (GBD) on WASP thereby displacing the VCA domain. While the V-verpolin-like motif binds actin monomer (G-actin), C-central and A-acidic domains bind and activate the ARP2/3 complex. Conformational changes induced by the binding of the ARP2/3 complex promote its binding to the actin filament, which is strengthened by the additional interaction of the ARP2/3 complex with WASP (VCA)-G-actin. Further conformational changes will secure the ARP2/3 complex on the filament and allow its binding to the actin monomer and the polymerization of the newly nucleated filament. Actin polymerizes at the fast-growing/barbed end, elongating towards the plasma membrane and the ARP2/3 complex would cross-link newly polymerizing filament to the existing filament. **(B)** ERK3 co-precipitates with active Rac1 and Cdc42 in complex with ARP2/3. Active Rac1/Cdc42 pull-down was performed using control and ERK3 knockdown HMECs. Levels of the active Rac1 and Cdc42 were assessed as well as the co-immunoprecipitation levels of ERK3, ARP2, ARP3, and ARPC1A. Levels of the total protein expression were evaluated in the total cell lysates (TCL) and Ponceau S staining was used as a loading control. **(C-F)** ERK3 regulates F-actin levels *in vitro* and *in vivo*. **(C)** Western Blot analyses of control (CRISPR Co) and ERK3-depleted (CRISPR ERK3) HMECs are presented alongside with representative confocal images of F-actin staining. **(D-E)** *In vivo* analysis of F- and G-actin levels in HMECs upon ERK3 knockdown. **(D)** Representative Western Blot analyses of the enriched F- and G-actin fractions as well as the ERK3 knockdown validation and total actin levels in the total cell lysates (TCL) are presented. **(E)** F- and G-actin levels were quantified, and ratios were calculated from five (n=5) independent experiments and are presented as mean ± SEM; *p<0.0332, **p<0.0021, ***p<0.0002, ****p<0.0001, unpaired t-test. Analyses of ERK3-dependent regulation of F-actin levels in cancerous MDA-MB231 cells is presented in Figure 5-figure supplement 1. Cellular colocalization between endogenous ERK3 and the ARP2/3 was assessed in the absence of Cdc42 and is presented in Figure 5-figure supplement 2. **(F)** Effect of full-length ERK3 on ARP2/3-dependent pyrene actin polymerization was assessed using a pyrene actin polymerization assay. Polymerization induced by the VCA domain of WASP which served as a positive control (green) as well as the ARP2/3 (orange) and ERK3 protein alone (blue) are shown for reference. Actin alone (black) was used to establish a baseline of polymerization. Fluorescence at 360/415 was measured over time and is presented as mean fold change from at least three independent experiments after normalization to the first time point within the respective group. ARP2/3-dependent actin polymerization was measured in the presence of both, ERK3 and WASP (VCA) domain and the results are depicted in Figure 5-figure supplement 3.

As ERK3 protein co-precipitated with active Rac1 and Cdc42 (Figure 2B and 2C) we further investigated whether also components of the ARP2/3 complex precipitate with Rac1/Cdc42. Interestingly, we readily detected the ARP2/3 complex subunits ARP3, ARP2 and ARPC1A as well as ERK3 by immunoblots in active Rac1/Cdc42 pull-downs (Figure 5B). Consistent with the reduction in active Rac1 and Cdc42 levels, we observed that knockdown of ERK3 by CRISPR/Cas9 or with si/shRNAs led to a reduction of F-actin staining in mammary epithelial cells as shown in Figure 5C and Figure 5-figure supplement 1A. To further quantify these effects and to corroborate the role of ERK3 in polymerization of actin filaments, we evaluated the ratio of cytoskeleton-incorporated filamentous (F-) and globular/monomeric actin (G-actin) in control and ERK3-depleted mammary epithelial cells by ultracentrifugation. Knockdown of ERK3 significantly shifted total F-actin to G-actin in both primary HMECs (Figure 5D and 5E) and MDA-MB231 cancer cells (Figure 5-figure supplement 1B and 1C). Moreover, in the absence of Cdc42, we could still detect colocalization between endogenous ERK3 and the ARP3 subunit of the ARP2/3 complex (Figure 5-figure supplement 2A-2D). These data prompted us to test whether ERK3 directly controls ARP2/3-dependent actin polymerization in the absence of Rac1 and Cdc42.

*In vitro,* the ARP2/3 complex expresses low intrinsic actin nucleating activity. *In vivo,* its activity is tightly controlled by NPFs which stimulate conformational changes^73, 74^. To determine whether ERK3 had any direct effect on the assembly of actin filaments, we used a pyrene-actin polymerization assay to monitor fluorescence kinetics over time as the actin filaments assembled in the presence of the purified ARP2/3 complex. We used the VCA domain of WASP protein as a positive control. Indeed, recombinant ERK3 stimulated ARP2/3-dependent actin polymerization at a nanomolar concentration (5 nM) in the presence of 10 nM of the ARP2/3 complex (Figure 5F). Moreover, the nucleation-promoting activity of ERK3 was comparable to WASP (VCA), a well-known NPF. ERK3 protein alone did not exert any stimulatory effect on actin nucleation (Figure 5F). These data suggest that ERK3 could function as nucleation promoting factor to promote ARP2/3-dependent actin polymerization. Additionally, we tested what effect it had on ARP2/3-dependent actin polymerization when both ERK3 and WASP (VCA) were present. Using the same concentration as for the initial screening (ERK3 4.8 nM and WASP (VCA) at 400 nM), we did not detect any additive effect when both proteins were combined (Figure 5-figure supplement 3). Combination of both proteins under this stoichiometric setting rather negatively affected the polymerization rate that could be achieved by each protein separately (Figure 5-figure supplement 3).

### ERK3 directly binds to ARP3

The ARP2/3 complex is composed of seven subunits: the actin-related proteins ARP2 and ARP3 and five associated proteins (ARPC1-5) that sequester ARP2 and ARP3 in the complex’s inactive conformation^35, 38^ (Figure 6A). *In vitro*, NPFs such as WASP stimulate actin polymerization by directly binding to and activating ARP2/3. Considering that ERK3 exerted a similar effect on the ARP2/3-dependent actin polymerization as NPFs, we further investigated the mode of interaction between ERK3 and the ARP2/3 protein complex by employing purified components. Indeed, using quantitative ELISAs we found that full-length ERK3 bound to the ARP2/3 complex with high affinity *in vitro* (Figure 6B).

**Figure 6.**
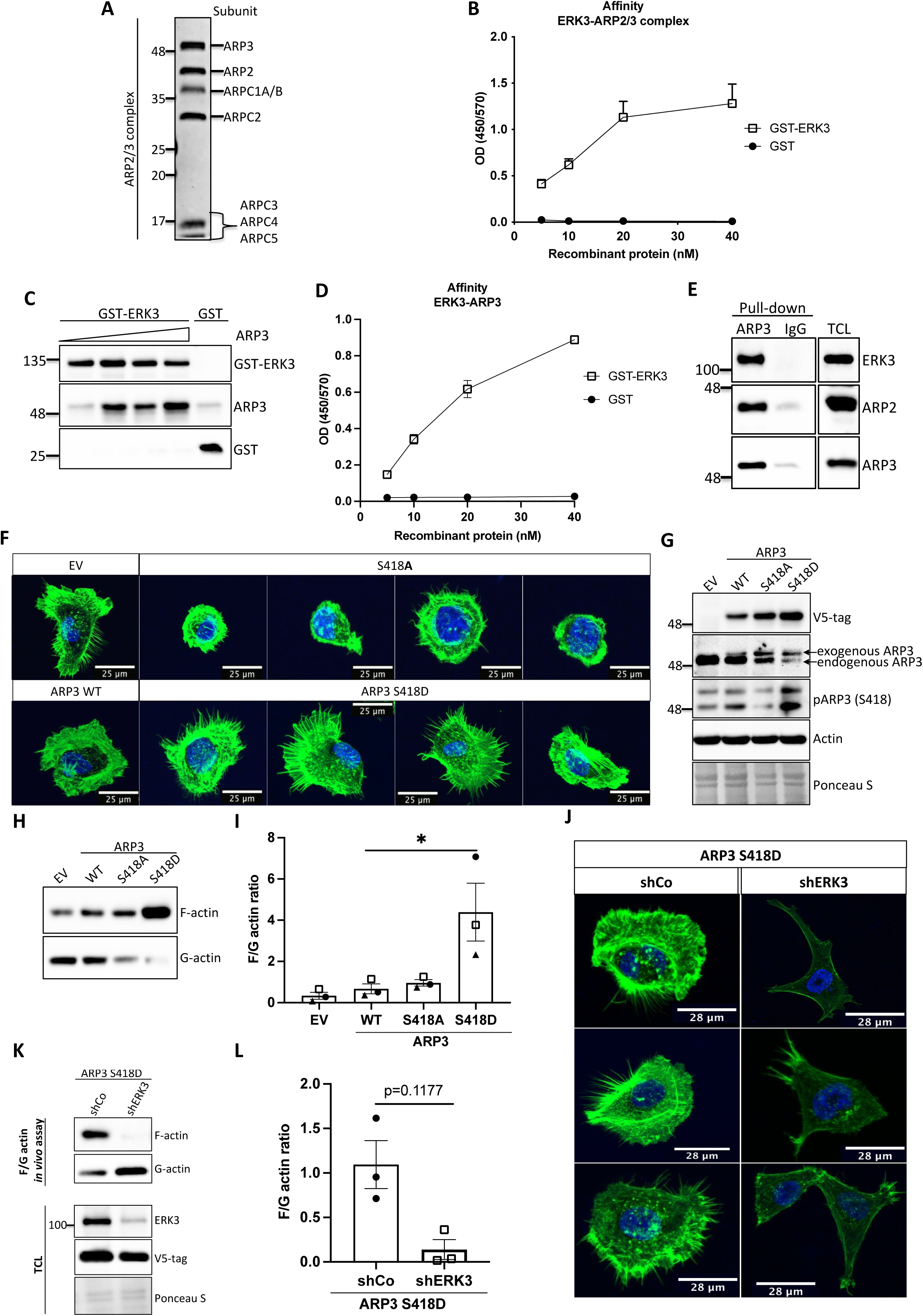
ERK3-dependent regulation of the ARP2/3 complex. **(A)** Coomassie stained 10% SDS-Page gel with 1 mg of the ARP2/3 protein complex (Cytoskeleton) presenting all the subunits. **(B)** Binding of increasing concentrations of recombinant GST-ERK3 to the ARP2/3 complex was measured by ELISA as described in the methods section. **(C)** The interaction between GST-ERK3 and ARP3 was measured *in vitro* using GST-pull-down assay as described in the methods section. **(D)** Binding affinity of the recombinant GST-ERK3 protein and ARP3 was assessed by ELISA as described in the methods section and mean absorbance (Abs) ± SEM from three independent experiments is presented. **(E)** Co-immunoprecipitation (IP) of ARP2/3 protein complex and ERK3 was performed in HMECs using ARP3 antibody. Levels of precipitated ARP3 as well as co-IP of ARP2 and ERK3 were assessed. IgG control was included to determine specificity of the interaction. Total cell lysate (TCL) was included to present expression levels of the verified interacting partners. Ponceau S staining was used as a loading control. **(F-G)** Actin phenotype of the HMECs was validated upon stable overexpression of the ARP3 non-phosphorylatable (S418A) and the phospho-mimicking (S418D) mutant, respectively. Wild type (WT) ARP3 was used as a control for the mutants and empty vector (EV) served negative control for the overexpression itself. **(F)** F-actin expression and organization in the negative (S418A) and phospho-mimicking (S418D) ARP3 mutant was visualized by green phalloidin and merged with the Hoechst staining of the nuclei. Four representative confocal images are presented. Images of EV-transfected and ARP3 WT-overexpressing HMECs are presented as controls. **(G)** Western Blot validation of the overexpression efficiency and phosphorylation of ARP3 at S418. Anti-V5-tag antibody was used to detect levels of exogenous ARP3 WT, S418A and S418D. Expression levels of the endogenous ARP3 were assessed as well as the phosphorylation at S418, total actin was validated. Ponceau S staining was used as a loading control. **(H-I)** Effect of the ARP3 mutant overexpression on F-actin levels was quantified using F/G actin *in vivo* assay. **(H)** Representative Western Blot analyses of F- and G-actin levels detected in fractions obtained from EV, ARP3 WT, S418A, S418D HMECs. **(I)** Quantification of the F/G actin ratios was performed for three (n=3) independent experiments and is presented as mean ± SEM; *p<0.0332, **p<0.0021, ***p<0.0002, ****p<0.0001, one-way ANOVA, Tukey’s post-test. **(J-L)** Effect of ERK3 depletion on dense F-actin phenotype of the ARP3 S418D- overexpressing HMECs. HMECs stably overexpressing ARP3 S418D were transduced with lentiviral particles targeting ERK3 (shERK3) and stable knockdown was established as described in the methods section. Cells were further subjected to analyses of the F-actin levels. **(J)** IF staining with OregonGreen Phalloidin 488 to visualize F-actin levels and organization. Scale bars 28 µm. **(K-L)** Effect of the ERK3 knockdown on F-actin levels was quantified in the ARP3 S418D mutant overexpressing HMECs using F/G actin *in vivo* assay. **(K)** Representative Western Blot analyses of F/G actin levels. ARP3 S418D-(V5-tagged) overexpression and ERK3 knockdown efficiency were validated in TCL. Actin and Ponceau S staining were used as loading controls. **(L)** Calculated ratios of F/G actin are presented as mean ± SEM from three (n=3) independent experiments; *p<0.0332, **p<0.0021, ***p<0.0002, ****p<0.0001, paired t-test. Colocalization of endogenous ERK3 with endogenous and exogenous ARP3 mutant (S418D) was verified and further effect of the ERK3 depletion on the Rac1 and Cdc42 activity was assessed in ARP3 S418D- overexpressing HMECs and presented in Figure 6-figure supplement 2.

ARP2 and ARP3 are essential for the nucleating activity of the ARP2/3 complex and occupy the central position in actin nucleation. Structural modeling has suggested that although ARP2 and ARP3 interact with each other in the inactive conformation, they are not not in the right configuration to act as a seed for actin nucleation. After activation by NPF binding the heterodimer mimics two actin subunits. Binding of an actin monomer then initiates the formation of the actin nucleus and the polymerization process^73, 75, 76^. Extensive characterization of the individual subunits also revealed that ARP3 is crucial for the stimulation and activity of the ARP2/3 complex and nucleation *in vitro*^75^. Considering that ARP3 is a structural component regulating nucleating properties of the ARP2/3 complex, we further determined whether ERK3 can directly bind to ARP3. Through GST-pull-down experiments with purified recombinant proteins, we could indeed detect a concentration-dependent direct binding between full-length ERK3 and ARP3 (Figure 6C). These results were further confirmed with ELISA analyses suggesting a high affinity binding between these two proteins (Figure 6D). Finally, we also detected ERK3 co-precipitating with ARP3 and ARP2 in human mammary epithelial cells at endogenous levels (Figure 6E).

### ERK3 phosphorylates ARP3 at S418

Conformational changes activating ARP2/3 are induced by binding to NPFs, actin and ATP. Additionally, phosphorylation of the ARP2/3 residues has been reported previously to play a crucial role in the conformational rearrangements within the ARP2/3 complex^77–81^. Most of the studies focused on phosphorylation of the ARP2 subunit, which induced the conformational repositioning of this subunit towards ARP3, further allowing binding of NPFs and full-activation of the ARP2/3 complex leading to the formation of the ARP2-ARP3 heterodimer^79–81^. From the seven components of the ARP2/3 complex ARP3, ARP2 and ARPC1A purified from *Acanthamoeba castellani* were shown to be phosphorylated^80^. Since ERK3 is a kinase, we assessed phosphorylation of the ARP2/3 complex by ERK3 by employing *in vitro* kinase assays and subsequent analysis of phosphopeptides by mass spectrometry. Initial analyses detected phosphorylation of the ARP3 subunit at serine 418 (S418) (peptide sequence: HNPVFGVM**S,** please see supplementary file 1). We further validated the significance of the detected phosphorylation *in vivo* by overexpressing the non-phosphorylatable (S418A) and phospho-mimicking (S418D) mutants of ARP3 in primary mammary epithelial cells and analysing the cell morphology and F-actin abundance. Cells transduced with wild-type (WT) ARP3 or empty vector (EV) were used as reference controls (Figure 6F). We were able to validate the phosphorylation of the ARP3 at Ser418 in the the wild type (EV) as well as the ARP3-overexpressing cells (Figure 6G) and detect a decrease at the S418 phosphorylation in ERK3-depleted cells depicted in Figure 5C (Figure 6-figure supplement 1). Of note, we observed that expression of exogenous ARP3 reduced the protein levels of the endogenous ARP3 (Figure 6G), possibly due to disruption of the protein complex, thereby affecting the stability of the endogenous proteins. Strikingly, expression of the phosphorylation-mimicking ARP3 mutant (S418D) under these settings predominantly induced actin filament formation, as indicated by the intense phalloidin staining (Figure 6F) and enhanced the F/G-actin ratio in primary mammary epithelial cells (Figure 6H and 6I). Moreover, S418D-overexpressing cells exhibited F-actin-rich protrusions (Figure 6F, lower panel). In contrast, most of the S418A-expressing cells were smaller in size and had a round morphology (Figure 6F, upper panel). Quantification of the F-actin levels revealed that despite the morphological distortion, cells expressing non-phosphorylatable ARP3 (ARP3 S418A) had no significant effect on overall F-actin levels (Figure 6H and 6I). To further determine the relevance of the S418 phosphorylation in ERK3-regulated ARP2/3-dependent actin polymerization in cells, we depleted ERK3 in the ARP3 S418D-overexpressing HMECs using shRNA (Figure 6J-6L). Knockdown of ERK3 led to a significant reduction of the F-actin levels in the ARP3 S418D-overexpressing cells. These data suggested that although phosphorylation at Ser418 of ARP3 promoted actin polymerization and thus the F-actin content it is not absolutely essential like ERK3. A concomitant active Rac1/Cdc42 pull-down revealed a significant decrease in Rac1 activity, with almost no effect on Cdc42 (Figure 6-figure supplement 2A and 2B). We could still detect ARP3 and ARP2 in complex with Cdc42 under these settings (Figure 6-figure supplement 2A). Using immunofluorescence analysis, we observed significant colocalization between endogenous ERK3 and ARP3 in HMECs (Figure 6-figure supplement 2C) and between endogenous ERK3 and overexpressed ARP3 S418D (Figure 6-figure supplement 2D-2F), respectively. Further, depletion of ERK3, as expected, reduced the migration of ARP3 S418D-expressing cells (Figure 6-figure supplement 2G and 2H).

### Kinase activity of ERK3 is necessary for membrane protrusions in primary mammary epithelial cells

ERK3 controls both F-actin levels by regulation of the new filament assembly via ARP3-binding and its branching and bundling into the actin-rich protrusions by binding to Cdc42/Rac1. We further investigated whether kinase activity of ERK3 was required for the formation of actin-rich protrusions and F-actin levels in mammary epithelial cells. Control (shCo) and ERK3-depleted (shERK3, 3’UTR) HMECs were reconstituted with wildtype (WT) or kinase dead (KD) (K49A/K50A) ERK3, and empty vector (EV) was introduced as a control (Figure 7A). Knockdown of ERK3 led to a decrease in actin-rich protrusions and overall F-actin staining (Figure 7B) with concomitant decrease in F-actin abundance (Figure 7C and 7D). Complementation with ERK3 WT rescued overall F-actin levels and the protrusive phenotype of the HMECs (Figure 7B,D). Interestingly, kinase-dead ERK3 recovered cytoskeletal F-actin levels to the same extent as ERK3 WT (Figure 7C and 7D). However, although the abundance of F-actin is rescued upon kinase-dead ERK3 overexpression, cells form dense meshwork of actin membrane ruffles which do not protrude into filopodia but rather curved around the edges of the cell forming a tangled web (Figure 7B). These results indicate that kinase activity of ERK3 is not required for the polymerization of actin, but rather for the bundling and/or branching of the rapidly polymerizing actin filaments. To further corroborate these observations, the wild type (WT) and kinase-dead (KD) ERK3 proteins were expressed in rabbit reticulocyte lysate (RRL) system and then employed to stimulate ARP2/3-dependent actin polymerization *in vitro*. Kinase activity of ERK3 does not seeem to affect the ARP2/3-dependent actin polymerization *in vitro* (Figure 7-figure supplement 1). We performed active Rac1/Cdc42 pull-down from HMECs shERK3 (3’ UTR) cells reconstituted with either wild-type (WT) or the kinase dead (KD) ERK3 and found that the kinase activity of ERK3 was not required for interactions or activation of Cdc42. However, Rac1 activation was partially dependent on the kinase activity of ERK3 (Figure 7-figure supplement 2A-2E).

**Figure 7.**
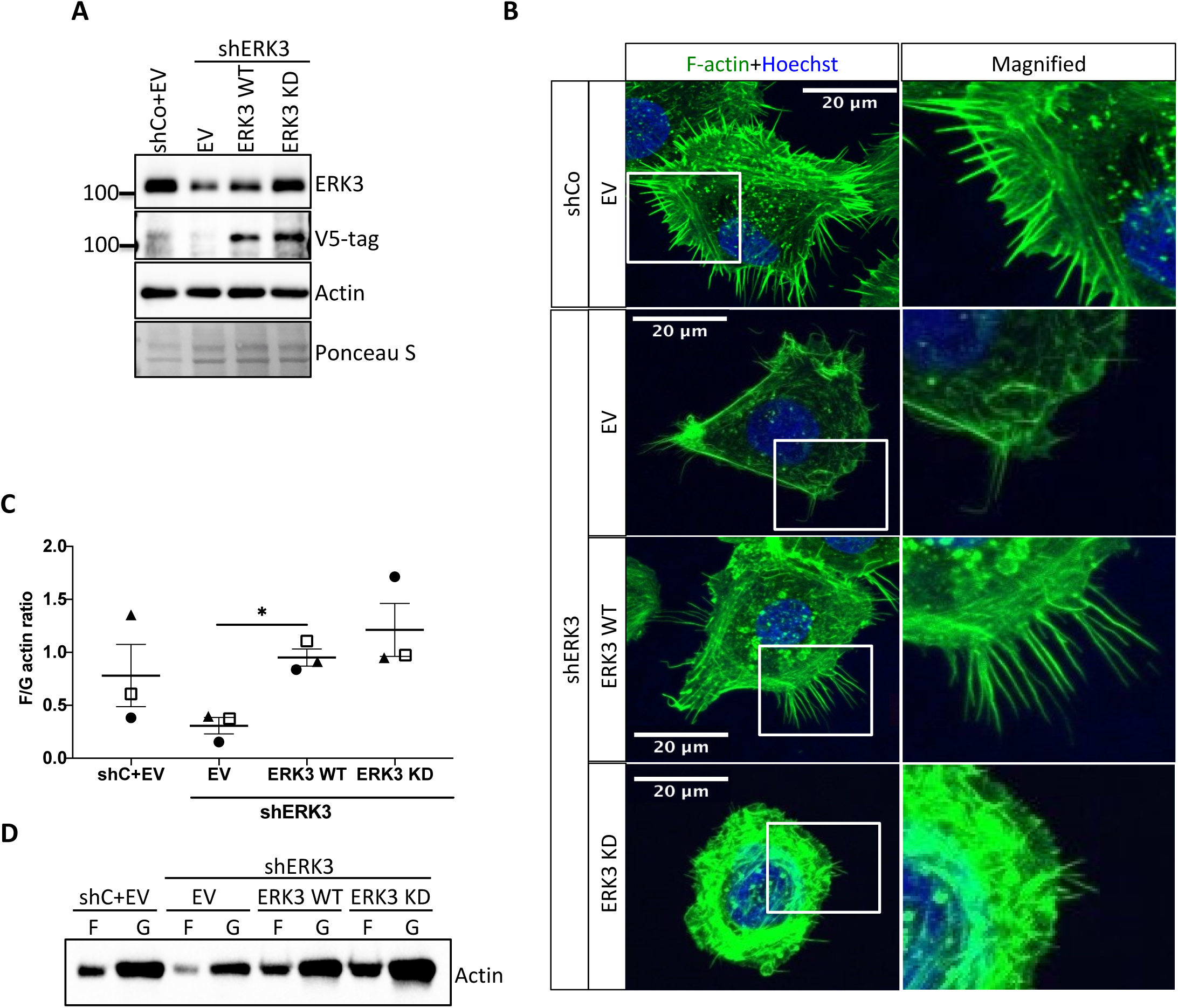
Kinase-dead mutant of ERK3 inhibits actin-rich protrusions with no negative effect on F-actin levels. **(A)** Western Blot validation of the ERK3 levels and expression of the V5-tagged ERK3 constructs is presented as well as the total actin levels. Ponceau S was used as a loading control. **(B)** Confocal analyses of control (shCo EV) and ERK3-depleted (shERK3 EV) HMECs reconstituted with wild type (ERK3 WT) or kinase-dead (ERK3 KD) (K49A/K50A) mutant of ERK3 after 4 h starvation. Merged images of F-actin (green, phalloidin) and Hoechst are shown on the left representing cell phenotype and magnified regions of the cell edges on the right show the actin distribution. **(C-D)** Levels of F-actin in HMECs analyzed by phalloidin staining were quantified using a F/G actin *in vivo* assay. **(C)** Calculated ratios of F/G actin are depicted as mean ± SEM from three (n=3) independent experiments; *p<0.0332, **p<0.0021, ***p<0.0002, ****p<0.0001, one-way ANOVA, Tukey’s post-test. **(D)** Representative Western Blot analyses of F- and G-actin levels.

**Figure 8.**
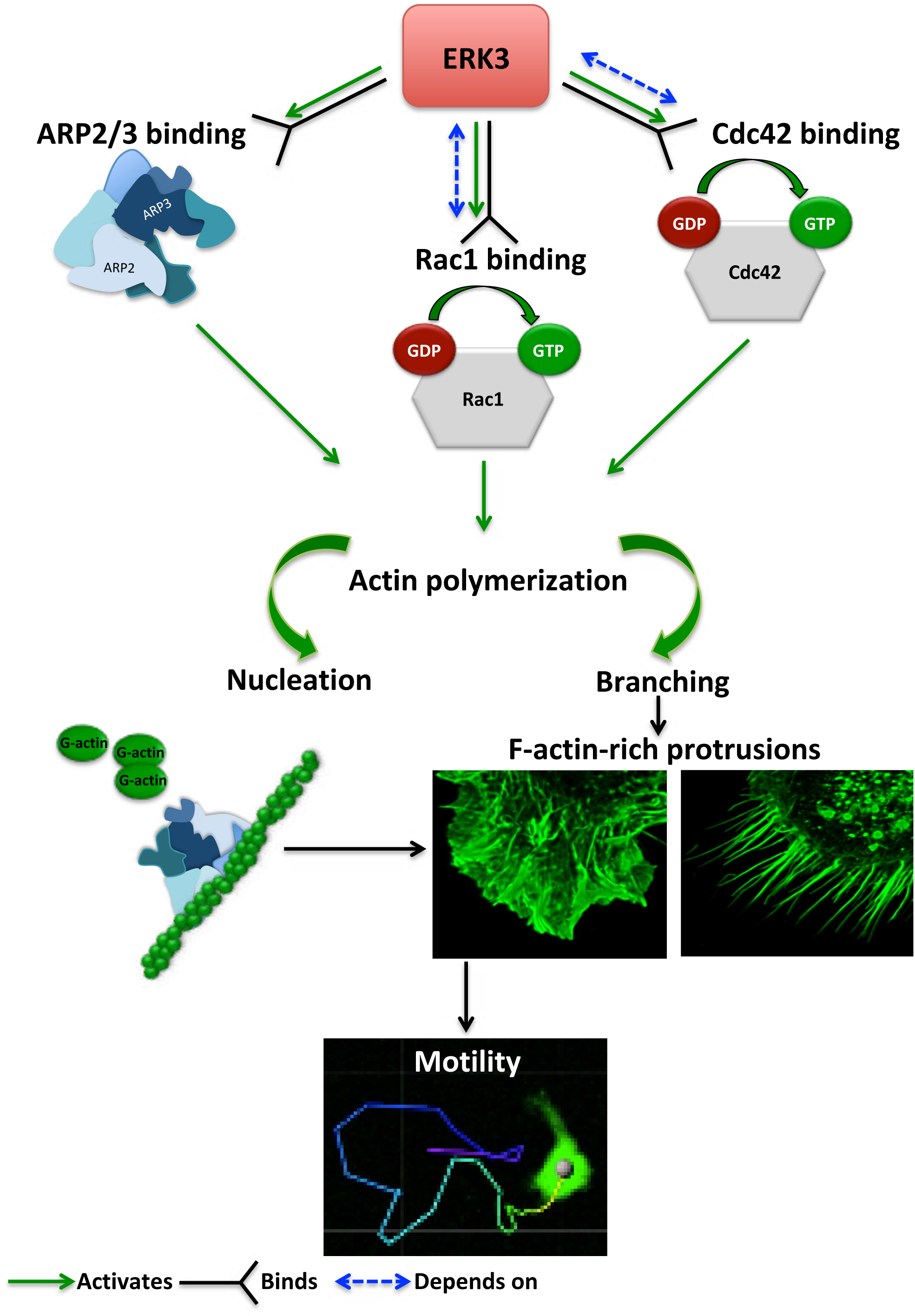
Schematic summary of the ERK3-dependent mechanisms regulating actin cytoskeleton and cell motility. ERK3 directly binds and activates ARP2/3 protein complex as well as the Cdc42 and Rac1 Rho GTPases. Activation of the ARP2/3 complex and Rac1/Cdc42 is required for nucleation of the new actin filaments, elongation and branching into the lamellipodia and filopodia. ERK3-regulates actin-rich protrusions which play a direct role in cell motility.

## Discussion

The regulation of the actin cytoskeleton is an intensely investigated area because of its fundamental role in many basic cellular functions. The signaling machinery that controls the actin skeleton dynamics is deregulated in cancer contributing to metastasis. Kinases are major drivers of signaling events and form the largest part of the druggable genome. Several drugs have successfully been developed to target deregulated kinases in cancer. Of the more than 500 kinases encoded by the human genome many studies have been focused on the conventional MAPKs, while the pathophysiological significance of atypical MAPKs remains underexplored. We have recently demonstrated that ERK3 directly contributed to AP-1/c-Jun activation, IL-8 production and chemotaxis^52^. Emerging studies have further shown that ERK3 functions in a context- and tissue-specific manner in controlling tumourigenesis and metastasis^50, 51, 53, 56, 81, 88, 89^. Mice expressing a catalytically inactive ERK3 mutant survive to adulthood and are fertile, but the kinase activity of ERK3 is necessary for optimal postnatal growth. In the present study, we aimed to elucidate how ERK3 influences cell shape and actin cytoskeleton and unexpectedly revealed a pivotal role for this atypical MAPK in the regulation of RhoGTPases as GEF, of ARP2/3- dependent polymerization of actin probably as NPF and of cell migration in mammalian cells. Further, our studies unveiled an evolutionarily conserved role for ERK3 in the formation of actin-rich protrusions.

### ERK3 is a GEF for Cdc42

The activation of both Rac1 and Cdc42 was compromised in the absence of ERK3 in primary mammary epithelial cells and oncogenic MDA-MB-231 cells. While the activation of Rac1 was partially rescued in human mammary epithelial cells upon EGF stimulation in ERK3-depleted cells, this was not the case in MDA-MB231 cells (Figure 2B, 2C and 2F, 2G). Furthermore, while ERK3 regulated the activity of both Rac1 and Cdc42 *in vivo* (Figure 2B-2G), the kinase domain of ERK3, which lacks kinase activity directly facilitated GDP-GTP nucleotide exchange specifically for Cdc42 (Figure 3B, 3C and 3G), which is implicated as a main regulator of filopodia assembly. In some cell types, we often detect a reduction in the total protein levels of Rac1 and Cdc42 after depletion of ERK3. Whether inactivation of these RhoGTPases prompt them for proteasomal degradation under these settings deserves further analysis. Using several experimental procedures, we demonstrated that interaction between ERK3 and Rac1 or Cdc42 is direct and nucleotide-independent which suggests that ERK3 is probably targeting the C-terminus of these RhoGTPases. Interestingly, it has been reported that Rac1 possesses a C-terminal docking (D site) for the canonical ERK (^183^KKRKRKCLLL^192^) ^45^. Whether the same site is being exploited by EKR3 for its binding to Rac1 or Cdc42 deserves further studies. It is interesting to note that while the kinase domain of ERK3 promoted GTP binding to Cdc42, it slightly attenuated the same process with Rac1 (Figure 3B-3G). While the kinase activity of ERK3 is not required for Cdc42 activation, Rac1 activity partially depends on it. Further structural, biochemical and biophysical studies are clearly warranted to unveil the molecular basis of this interaction and the subsequent functional implications.

In cells, regulation of RhoGTPases and that of the actin assembly is a much more complex and integrative process^68^. For example, a cell type-dependent, hierarchical crosstalk between RhoGTPases was observed in the regulation of actin-rich protrusions and Cdc42 activity was shown to initiate Rac1-dependent lamellipodia protrusions^67, 90, 91^. While ERK3 might activate Rac1 through its GEF-like function of Cdc42, it could also function as an intermediate link between Rac1 and its GEF by scaffolding both proteins and/or activating the GEF, thus indirectly contributing to Rac1 activity.

It is highly interesting that PAK family kinases, the effector kinases of Rac1 and Cdc42, directly phosphorylate the SEG motif in the activation loop of ERK3^60, 61^. The observations presented here strongly suggest a positive feed-back loop, where ERK3 is required for the GTP loading of Cdc42 and Rac1 to induce their interaction with PAKs which in turn phosphorylate and activate ERK3 and we show that active ERK3 kinase is required for eventual formation of actin-regulated membrane protrusions. The functional and physical uncoupling of this loop could possibly be controlled by a phosphatase or by inducing the rapid turn-over of ERK3 protein. The constituents of this dynamic complex need further evaluation and characterization.

### ERK3 binds ARP3 to regulate its function in actin polymerization

Cells rely on several mechanisms to regulate the assembly of actin filaments. To complicate matters further, regulatory pathways involved in the filopodia formation seem to be cell-type specific and Cdc42-, WASP- and ARP2/3-independent mechanisms of membrane spikes formation have been proposed^92–94^. In this study, we demonstrate that ERK3 directly binds the ARP2/3 complex and contributes to actin polymerization (Figure 5 and 6). Conformational changes in ARP2/3 complex assembly are crucial for the initial heterodimer formation between ARP2 and ARP3 subunits^42, 75, 95^. In the unstimulated complex, ARP2 and ARP3 subunits exist in the splayed confromation and upon stimulation by NPFs the two proteins align into a side-by-side position creating a surface mimicking the first two subunits of the new actin filament ^35, 96^. Binding of these two crucial subunits by NPFs could reassure an active conformation. Interestingly, phosphorylation of certain subunits of ARP2/3 complex has been shown to be indispensable for the destabilization of its inactive conformation prior the full activation by the NPFs^77, 79–81^. LeClaire et al., further reported that Nck-interacting kinase (NIK) regulates ARP2 phosphorylation and grooms ARP2/3 complex for the NPFs activation. Phosphorylation events within the ARP2/3 complex could regulate nucleation activity by affecting several properties, such as complex formation itself, conformational changes in the complex assembly or the affinity of the ARP2/3 to NPFs. Further structural simulation studies are needed to test if the interaction of ERK3 with ARP2/3 complex and the ARP3 phosphorylation at S418 inflict key conformation changes to support actin polymerization. While we uncover a role for ERK3 in the formation of filopodia and ARP2/3-dependent actin polymerization, the possible role for ERK3 in the regulation of other actin nucleation factors including formins cannot be ruled out.

While the kinase activity of ERK3 is clearly required for the formation of actin-rich protrusions, expression of kinase-dead constructs in ERK3-depleted cells failed to inhibit F-actin enrichment. These data suggest that perhaps ERK3 binding to ARP2/3 and possibly other hitherto unknown factors per se contributes to actin polymerization, while its kinase activity is probably still required for ARP2/3- dependent and Cdc42-induced filopodia formation, polarization and migration of mammalian epithelial cells.

Overall, our studies shed new light on the multi-layered regulation of actin cytoskeleton by an understudied MAPK. ERK3 is druggable and kinase inhibitors have already been developed for clinical use, which could also serve as a tool to decipher kinase-dependent and -independent functions. These observations will not only enhance the current understanding of actin cytoskeleton regulation, but also instigate new lines of investigations on the molecular, structural and functional characterization of RhoGTPases and the ARP2/3 complex.

## Material and methods

### Cell culture

Immortalized Human mammary epithelial cells (HMECs) hTERT-HME1 (ME16C) (ATCC CRL-4010) were purchased from ATCC (Manassas, VA 20108 USA) and cultured in mammary epithelial cell growth medium (MEGM) BulletKit, containing supplements and growth factors (Cat# CC-3150, Lonza) according to the Clonetics^®^ mammary epithelial cell system quide (Lonza). Triple-negative breast cancer cell line MDA-MB231 was purchased from DSMZ (DSMZ# ACC 732) and cultured in Dulbecco’s Modified Eagle Medium (DMEM) supplemented with 10% heat inactivated FBS. MDA-MB231 cells overexpressing GFP actin were a kind gift from Prof. Dr. Mohamed Bentires-Alj (University of Basel, department of Biomedicine). 293T cells were a kind gift from Dr. Andreas Ernst (Goethe-University Frankfurt am Main, IBC2) and were cultured in DMEM supplemented with 10% FBS. MDA-MB231 cells overexpressing GFP were a kind gift from Prof. Dr. Mohamed Bentires-Alj (University of Basel, department of Biomedicine). MDA-MB468 cells were purchased from DSMZ and cultured in DMEM/F-12 medium (Cat# 11320033, Gibco) supplemented with 10% FBS. MCF7 cells were cultured in Roswell Park Memorial Institute 1640 (RPMI 1640) medium (Cat# 11875093, Gibco) supplemented with 10% FBS, 1% MEM non-essential amino acids (Cat# M7145, Sigma), 1mM sodium pyruvate and 10 µg/ml human insulin (Cat# 19278, Sigma). HUVECs were a kind gift from Prof. Dr. Wolfram Ruf (Center for Thrombosis and Hemostasis (CTH), University Medical Center of the Johannes Gutenberg University Mainz) and were cultured on 0.2% gelatin coating in Endothelial Cell Growth Medium (Cat# C-22010, PromoCell). HT-29 cells were purchased from ATCC (ATCC HTB-38) and cultured in McCoy’s medium supplemented with 10% heat inactivated FBS. Calu-1 cells were obtained from Sigma and cultured in Dulbecco’s Modified Eagle Medium (DMEM) supplemented with 10% heat-inactivated FBS.

All cell lines used in this study were authenticated cell lines obtained from ATCC or DSMZ. All used cells were periodically tested for *Mycoplasma* contamination with negative results.

### Stimulation of cells

HMECs and MDA-MB231 cells were seeded in 12-well plates at initial density of 2×10^5^ cells/well. After cells reached 70% confluence, medium was exchanged to MEGM with no supplements for HMECs and DMEM minus FBS for MDA-MB231 cells, 4h prior treatment with recombinant human epidermal growth factor (EGF) (Cat# RP-10927, Invitrogen) at 5, 10, 15 and 30 minutes. Afterwards, cells were subjected to Western Blot analyses.

### Cycloheximide (CHX) chase experiments

To determine ERK3 half-life in mammary epithelial cells and its alteration upon EGF treatment, HMECs were seeded 12-well plates at an initial density of 2×10^5^ cells/well and cultured until 70% confluent. Medium was exchanged to MEGM with no supplements, 4 h before cells were treated with human recombinant EGF at 100µg/ml for 15 min, followed by treatment with protein synthesis inhibitor cycloheximide (CHX) (Cat# C-7698, Sigma, stock 100 mg/ml in DMSO) at 0.5, 1, 2, 4 and 6 h. Cells were subjected to Western Blot analyses and quantification. Fold change in ERK3 protein levels was calculated in respect to the untreated cells (-EGF, 0 h) or to the respective control in each group (0 h) for unstimulated (-EGF) and EGF-stimulated (+EGF) cells using ImageJ software.

### Antibodies

Primary antibodies used: phospho-ERK3 (pSer189) (Cat# SAB4504175, Sigma), ARPC1A (Cat# HPA004334, Sigma), ERK3 (Cat #4067, Cell Signaling Technology (CST)), phospho-p44/42 MAPK (Thr202/Tyr204) (Cat# 9101L, CST), V5-tag antibody (Cat # R960-25, Invitrogen), GST antibody (Cat# 2622S, CST), normal rabbit IgG antibody (Cat# 2729, CST), GST (B-14) antibody (Cat# sc-138, Santa Cruz Biotechnology), Rac1 (Cat# 610651, BD Transduction Laboratories), Cdc42 (Cat# 610929, BD Transduction Laboratories), ARP3 (phospho-Ser418) antibody (Cat# orb317559) was obtained from Biorbyt and its specificity was validated in cells overexpressing active (S418D) and phoshpo-impaired (S418A) ARP3 mutants (Figure 6G). ARP3 (Cat# ab151729, Abcam), ARP2 (Cat# ab128934, Abcam), beta-actin HRP conjugated antibody (Cat# ab49900, Abcam). HRP-conjugated secondary antibodies for rabbit and mouse IgG were obtained from Invitrogen (Cat# A16096 and A16066, respectively).

Antibodies used for the IF staining: ERK3 antibody (Cat# MAB3196, R&D).

### Western blotting

Cells were washed with ice-cold PBS (10 mM sodium phosphate, 150 mM NaCl, pH 7.2) and lysed in cold RIPA lysis buffer: 250mM NaCl, 50mM Tris (pH 7.5), 10% glycerol, 1% Triton X-100), supplemented with protease inhibitor cocktail Set I-Calbiochem 1:100 (Cat# 539131, Merck Millipore) and phosphatase inhibitors: 1 mM sodium orthovanadate (Na_3_VO_4_), 1 mM sodium fluoride (NaF). Cells were lysed for 30 min on ice, followed by 10 min centrifugation at 14000 rpm. Protein concentrations were estimated using 660 nm Protein Assay (Cat# 22660, ThermoFisher Scientific). Samples were prepared by mixing with 4 x SDS-PAGE sample buffer (277.8 mM Tris-HCl pH 6.8; 44.4% glycerol, 4.4% SDS, 0.02% bromophenol blue) supplemented with 50 mM dithiothreitol (DTT) and boiling at 95°C for 5 min. Samples were then subjected to SDS-PAGE followed by transfer of the proteins onto nitrocellulose membranes (GE Healthcare, Chalfont St Giles, UK). Membranes were blocked in 3% BSA/PBST (1x PBS, pH 7.5 containing 0.5% Tween-20) for 1 h at RT and incubated with primary antibody diluted in PBST at 4°C, overnight. Following 3 x 5 min washing with PBST membranes were incubated with HRP-conjugated secondary antibody for 1 h at RT. After washing, signal was visualized using chemiluminescent HRP substrate (Immobilon Western, Cat# WBKLS0500, Merck Millipore). Western Blots semi-quantification was performed using ImageJ software (ImageJ, RRID: SCR_003070).

### Site-directed mutagenesis and expression constructs

Full-length ERK3 wild type (WT) pDONR-223 construct purified from Human Kinase Library (Addgene) was used as a template to generate ERK3 K49A K50A kinase dead (KD) mutant. Mutations were introduced using the following primers were used: frw_5’ GCAATTGTCCTTACTGATCCCCAGAGTGTC, rev_5’ CGCGATGGCTACTCTTTTGTCACAGTC using Q5 High-Fidelity DNA Polymerase (Cat# MO491, New England BioLabs). For lentiviral expression, ERK3 WT and ERK3 K49A K50A (KD) mutant were cloned into pLenti4TO/V5-Dest expression vector.

Wild type ARP3 pENTR223 construct (Harvard HsCD00375598) was cloned into pLenti4TO/V5-Dest mammalian expression vector and pGEX-6-P1 for bacterial expression. Serine 418 (S418) phopsho-dead (S418A) and phosphor-mimicking (S418D) constructs of ARP3 were obtained from Eurofins (Cat# #11104144057-1-5 and #11104144057-1-6, respectively) and cloned into the pLenti4TO/V5-Dest expression vector.

### Generation of knockdowns

shRNAs plasmids were purified from the MISSION® shRNA Human Library (Sigma). Non-targeting control shRNA (shCo) MISSION® pLKO.1 puro (Cat# SHC001) was included.

shMAPK6 (shERK3) TRCN0000001568, NM_002748.x-3734s1c1,

CCGGGCTGTCCACGTACTTAATTTACTCGAGTAAATTAAGTACGTGGACAGCTTTTT

shCdc42#1 TRCN0000047628, NM_001791.2-471s1c1

CCGGCCCTCTACTATTGAGAAACTTCTCGAGAAGTTTCTCAATAGTAGAG GGTTTTTG

shCdc42#2 TRCN0000047629, NM_001791.2-193s1c1

CCGGCGGAATATGTACCGACTGTTTCTCGAGAAACAGTCGGTACATATTC CGTTTTTG

siRNAs were purchased from Qiagen:

siMAPK6#1 (siERK3#1) FlexiTube siRNA 5 nmol, siRNA Name: Hs_MAPK6_5, Cat# SI00606025, sense strand: 5’-AGUUCAAUUUGAAAGGAAATT-3’. Negative control siRNA (siCo) Cat# 1027310.

### siRNA transfection

Cells were seeded one day before transfection at an initial density of 2×10^5^ cells/well in 12-well plates or at 3×10^5^ cells/well (6-well plates). Cells were transfected using SAINT-sRNA transfection reagent (SR-2003, Synvolux) according to the manufacturer’s instructions. Cells were analysed 48 h post-transfection and knockdown efficiency was verified by Western Blot using target-specific antibodies. If EGF stimulation was included in the experimental settings, 48 h post-transfection medium was exchanged for serum/supplements-free medium for 4 h prior the EGF (100 ng/ml) treatment for 15 min.

### Lentiviral-mediated knockdown and expression of cDNAs

For the production of lentiviral particles, the following packaging plasmids were used: pHDM-G (encoding VSV-G), pHDM Hgpm2 (encoding codon-optimized HIV gag-pol proteins), pHDM tat 1b (encoding HIV Tat1b protein) and pRC CMV-Rev1b (encoding HIV rev protein). Lentiviral particles were generated following standard protocols. Briefly, lentiviral supernatants were produced in 293T cells by co-transfection of the cells with lentiviral packaging plasmids (0.3 µg each), lentiviral expression constructs (1 µg) and 10.8 µl of 10 mM polyethylenimine (PEI). The viral particles were harvested after 48 h and sterile-filtered. Cells were infected with lentiviral particles in the presence of 10 µg/ml of polybrene (Cat# sc-134220, Santa Cruz) and selected with puromycin at 3 µg/ml (Cat# 0240.3, Carl Roth).

For complementation assays, empty vector control (pLenti4TO/V5-Dest), WT or KD (K49A K50A) mutant of ERK3 were reintroduced into shERK3 (3’UTR) background by lentiviral transduction and cells were double-selected with zeocin (100 µg/ml) (Cat# R25001, Invitrogen) and puromycin.

To generate CRISPR/Cas9 mediated ERK3 knockout in the HMECs, cells were infected with lentiviral particles and selected with puromycin (8 µg/ml). Lentiviral particles coding for CRISPR ERK3 and CRISPR control vector (pLentiCRISPRv2) were produced in 293T cells by co-transfection of lentiviral packaging plasmids (0.3 µg each) and 1.1 µg of lentiviral vector containing the respective gRNAs in the presence of 21 µl of Lipofectamine2000 (Cat# 11668027, ThermoFisher Scientific). CRISPR/Cas gRNA sequences targeting ERK3 were designed by Rule Set 2 of Azimuth 2.0 as described previously ^97^. The top three scoring gRNAs were selected:

#1 5’-CACCGAGCCAATTAACAGACGATGT-3’

#2 5’-CACCGATACTTGTAACTACAAAACG-3’

#3 5’-CACCGCTGCTGTTAACCGATCCATG-3’

gRNAs were individually cloned into pLentiCRISPRv2 (Addgene plasmid #52961), following established protocols ^98^.

### Transient transfections

HMEC cells stably transfected with shRNA targeting ERK3 at 3’UTR (shERK3) or with control empty vector shRNA (shCo) were transiently transfected with either an empty vector pcDNA3/V5-Dest40 (EV), ERK3 WT or ERK3 K49A K50A, ERK3 S189A or S189D mutant construct (0.5 µg plasmid) in the presence of PEI (3 µl/well). 6 h post-transfection medium was exchanged for DMEM + FBS complete medium. 24 h post-transfection medium was exchanged again for DMEM–FBS medium. 48 h post-transfection supernatants were harvested for IL-8 ELISA and cells were analysed by Western Blot.

The images of both GFP fluorescence and transmission of the live cells were acquired with a Leica SP8 confocal microscope (Leica, Mannheim, Germany) was performed with a 10X 0.3 NA objective, with 488 nm excitation (at approximately 150 µW), with an emission window of 500 nm to 590 nm for GFP detection and with scanning Differential Interference Contrast transmission imaging in a 1500 µm x 1500 µm frame format with 400 lines per second, 0.71 µm/pixel (2048 x 2048 pixel per frame) and with 2 times averaging per line with a frame acquisition of every 30 minutes per selected position within the chamber.

Phosphorylation of the ARP2/3 protein complex subunits by ERK3 was detected by mass spectrometry analyses after in vitro kinase assay. Proteins were in-gel digested and subsequently enriched for phosphopeptides using TiO2 beads.

### RNA isolation, cDNA synthesis and RT-PCR analysis

For gene expression analyses, cells were washed with cold PBS and total RNA was extracted using Trizol (Cat# 15596018, Ambion) according to the manufacturer’s instructions. Isolated RNA was then used as a template for cDNA synthesis with the RevertAid First Strand cDNA synthesis kit (Cat# K1621, ThermoFisher Scientific) and random hexamer primers.

Real-time PCR was performed using EvaGreen qPCR master mix (5 x Hot Start Taq EvaGreen® qPCR Mix (No ROX), Cat# 27490, Axon) and following primers:

*ERK3* Frw_5’ ATGGATGAGCCAATTTCAAG

Rv_5’ CTGACAATCATGATACCTTTCC

The housekeeping gene for human 18S was used for normalization:

*18s* Frw_5’ AGAAACGGCTACCACATCCA

Rv_5’ CACCAGACTTGCCCTCCA;

Relative expression levels were calculated as ΔΔCt and results are presented as log2fold change in gene expression.

### Protein purification

The pGEX-2T-Rac1-WT and pGEX-2T-Cdc42-WT bacterial expression constructs were a gift from Gary Bokoch (Addgene plasmid# 12977 and #12969, respectively). The pGEX2TK-Pak1 (70-117) containing the p21-binding domain (PBD) was a gift from Jonathan Chernoff (Addgene plasmid# 12217). For the purification of ARP3 (ACTR3), construct (Harvard HsCD00375598) in pGEX-6-P1 expression vector was used.

Rosetta (DE) competent cells expressing Rac1-WT, Cdc42-WT or ARP3 were incubated at 37 °C, followed by induction of protein synthesis with 0.1 mM Isopropyl β-D-1-thiogalactopyranoside (IPTG) at 16 °C, overnight. Bacteria were lysed in 1% NP-40, 20 mM Tris–HCl, pH 7.5, 200 mM NaCl buffer. Sonicated lysates were used for the GST purification using PureCube glutathione agarose (Cat# 32105, Cube Biotech). After incubation, beads were washed with PBS pH 7.5.

Enzymatic cleavage of the GST tag was performed using thrombin (Cat# T1063, Sigma). Supernatants containing cleaved target proteins were incubated with Benzamidine Sepharose 4 flat flow (Cat# 17-5123-10, Cytvia) to remove thrombin. If required, eluted proteins were then concentrated using amicon ultra-15 centrifugal filters (Cat# UFC901024, Merck Millipore).

Proteins size and purity were assessed throughout the purification procedures.

### Synthesis of wild type (WT) and kinase dead (KD) ERK3 proteins using rabbit reticulocyte lysates (RRL)

ERK3 WT or ERK3 kinade dead (KD) (K49A K50A) in pcDNA3/V5-Dest40 vector were expressed in T7 rabbit reticulocyte lysates using TNT T7 Quick Coupled transcription/Transaltion System (Cat# TM045, Promega) following manufacturer’s protocol. Expression levels were assessed by Western Blot, using V5-tag specific antibody. Lysates were further used in actin polymerization assay as recombinant proteins.

### Endogenous pull-down of active Rac1/Cdc42

Levels of GTP-bound Rac1 and Cdc42 were determined using either active Rac1/Cdc42 pull-down and detection kit (Cat# 16118/19, ThermoFisher Scientific) following the of the manufacturer’s instructions and buffer composition or purified GST-Pak1-PBD fusion beads. HMEC and MDA-MB231 cells were seeded in 6-well plates at an initial density of 3×10^5^ cells/well. After cells reached 70% confluence, medium was exchanged to MEGM (no supplements) for HMEC or DMEM without FBS for MDA-MB231 cells, 4 h prior 15 min stimulation with 100 ng/ml of recombinant human EGF.

Afterwards, cells were subjected to active Rac1/Cdc42 pull-down. The precipitated samples and total cell lysates were subjected to Western Blot analyses and ImageJ quantification. Relative levels of active Rac1 and Cdc42 were determined by calculating the ratio of active (GTP loaded) RhoGTPases with respective total protein levels in TCL.

### Endogenous pull-down of ARP3

HMECs were were seeded in 6-well plate at an initial density of 2×10^5^ cells per well and cultured until 80% confluent. For the immunoprecipitation (IP) of the endogenous ARP3, cells were washed with ice-cold PBS and lysed with ice-cold IP buffer (10 mM HEPES pH 7.4; 150 mM NaCl, 1% Triton X-100, plus protease inhibitor cocktail Set I-Calbiochem 1:100 (Cat# 539131, Merck Millipore), 1 mM Na_3_VO_4_ and 1mM NaF). Cell lysates were incubated with Protein A/G-Agarose beads (Cat# 11 134 515 001/ 11 243 233 001, Roche), and either ARP3 or normal Rabbit IgG antibody for 2 h at 4°C with rotating. After the incubation beads were washed with IP buffer and analysed by immunoblot. Levels of the immunoprecipitated ARP3 as well as the co-immunoprecipitation of ARP2 and ERK3 were assessed. Total cell lysates were used as a control.

### Binding interaction-GST pull-down assays

Purified recombinant GST-fusion Rac1 and Cdc42 proteins immobilized on the beads were used for *in vitro* GST pull-down experiments.

To verify the relevance of the nucleotide binding in the interaction of the GTPases with ERK3 protein, Rac1 and Cdc42 proteins were loaded with non-hydrolysable GTPγS or GDP (Components of the active Rac1/Cdc42 pull-down and detection kit (Cat# 16118/19, ThermoFisher Scientific).

Recombinant GST-fusion GTPases protein beads were incubated with gentle shaking for 15 min at 30°C in 100µl of 25mM Tris-HCl, pH 7.2, 150mM NaCl, 5mM MgCl_2_, 1% NP-40 and 5% glycerol binding/wash buffer containing 0.1mM of GTPγS or 1mM GDP in the presence of 10mM EDTA pH 8.0 to facilitate nucleotide exchange. Reaction was terminated by placing samples on ice and addition of 60mM of MgCl_2_. GTPγS-/GDP-loaded Rac1 and Cdc42 beads were centrifuged and supernatants were removed. Beads were subjected to GST pull-down ERK3 binding assay.

GTPγS/GDP-loaded GSt-Rac1 and GST-Cdc42 protein beads were incubated with recombinant human ERK3 protein (Cat# OPCA01714, Aviva Systems Biology) in 100µl of binding/wash buffer supplemented with protease inhibitor with protease inhibitor cocktail Set I-Calbiochem 1:100 (Cat# 539131, Merck Millipore) and phosphatase inhibitors: 1 mM sodium orthovanadate (Na_3_VO_4_), 1 mM sodium fluoride (NaF) for 1h at 4°C with rotation. Recombinant GST protein bound to glutathione beads was used as a negative control. After the incubation, beads were washed 3 times with 400µl binding/wash buffer, centrifuged each time at 2000 rpm. Samples were eluted with 4x laemmli buffer supplemented with 100mM DTT and 5 min boiling at 95°. Samples were subjected to Western Blot analyses.

For the in vitro interaction between GST-Rac1/ GST-Cdc42 and ERK3 kinase domain, Rac1/Cdc42 WT beads and ERK3 (amino acids (aa) 9-327) (Crelux) E. coli purification was employed. Additional information on the Crelux protein can be viewed in the Supplementary information file 1.

For ERK3-ARP3 binding studies, human recombinant GST-fusion ERK3 protein (SignalChem) and E.coli purified ARP3. Glutathione beads were incubated with 37 nM of GST-ERK3 or GST alone and indicated concentrations of ARP3 for 2 h at 4°C in the binding buffer (5mM Tris-HCl, pH 8.0, 0.2mM CaCl_2_) supplemented with protease and phosphatases inhibitors. Afterwards, beads were washed 3 times with 400µl binding buffer supplemented with 1% NP-40. Samples were eluted with 4x laemmli buffer supplemented with 50 mM DTT and 5 min boiling at 95°. Immobilized protein complexes were detected by GST and ARP3 specific antibodies.

### Determination of GTP loading on Rho GTPases

#### *In vitro* active Rac1/Cdc42 pull-down assay

The efficiency of the nucleotide loading under chosen conditions of the GTP-loading status of the purified Rac1 and Cdc42 proteins were determined using GST-Pak1-PBD fusion beads.

Recombinant Rac1 and Cdc42 proteins (no tag) were loaded with GTPγS or GDP by incubation for 15 min at 30°C in 100µl of 25mM Tris-HCl, pH 7.2, 150mM NaCl, 5mM MgCl_2_, 1% NP-40 and 5% glycerol binding/wash buffer containing 0.1mM of GTPγS or 1mM GDP in the presence of 10mM EDTA pH 8.0. Reaction was terminated by placing samples on ice and addition of 60mM of MgCl_2_. GTPγS-/GDP- loaded Rac1 and Cdc42 as well as the native (WT) proteins were subjected to active Rac1/Cdc42 pull-down assay using GST-Pak1-PBD fusion beads. Levels of GTP loaded GTPases were detected using Rac1 or Cdc42 specific antibodies.

#### *In vitro* binding assay- recombinant protein ELISA

Binding affinity of ERK3 protein to Rac1, Cdc42, ARP2/3 complex and ARP3 protein was measured by enzyme-linked immunosorbent assay (ELISA). Purified Rac1, Cdc42 and ARP3 proteins or ARP2/3 protein complex (Cytoskeleton) were used as bait and full-length GST-ERK3 (SignalChem) was used as a titrant protein. Purified GST protein was used as a negative control for ERK3 binding. Each reaction was ran in triplicates. 96-well Nunc MaxiSorp^TM^ plates (Cat# 44-2404-21, ThermoFisher Scientific) were coated with 100 ng/well of bait protein diluted in 100 µl of PBS pH 7.5 overnight at 4°C. Plates were washed with 300 µl/well of 0.5% Tween-20/PBS pH 7.5 and blocked with 300 µl of 1% BSA in PBS pH 7.5 for 1 h at room temperature with gentle shaking. After washing with Tween-20/PBS, titrant proteins were added at 5, 10, 20 and 40 nM final concentration in 100 µl of 1% BSA/PBS and plates were incubated for 2 h at RT with gentle shaking. Afterwards, plates were washed with Tween-20/PBS and incubated with primary antibodies: anti-GST or anti-ERK3 specific antibody (1:500) in 100 µl/well of 1% BSA/PBS for 1 h at RT, followed by washing with Tween-20/PBS and 1 h incubation with HRP-conjugated goat anti-mouse IgG at 1:40 000 (Cat# A16066, Invitrogen) in 100 µl per well of Tween-20/PBS. After washing with Tween-20/PBS, plates were incubated with 100 ml/well of eBioschence^TM^ TMB substrate solution (Cat# 00-4201-56, Invitrogen). Reactions were stopped with 50 µl of STOP solution (1 M H_3_PO_4_). Absorbance was measured at 450 nm with 570 nm reference using microplate reader Tecan SPARK^®^. Multiple negative controls were used to establish each affinity assay, including: coating protein + 1^st^ Ab, coating protein+ 2^nd^ Ab, coating protein + 1^st^ Ab + 2^nd^ Ab, coating protein + titrant protein + 1^st^ Ab only, coating protein + titrant protein + 2^nd^ Ab only, coating protein + BSA + 1^st^ Ab + 2^nd^ Ab, coating with BSA + titrant protein + 1^st^ Ab + 2^nd^ Ab.

#### GDP/GTP nucleotide exchange assays

GDP/GTP-exchange assay was performed using recombinant Rac1 and Cdc42 proteins in binding buffer containing: 20 mM Tris-HCl (ph 7.5), 100 mM NaCl and protease/phosphatases inhibitors. Firstly, RhoGTPases were stripped of nucleotides by incubation with binding buffer containing 10 mM EDTA for 10 min at RT. Afterwards, reactions were supplemented with 50 mM MgCl_2_ and 500 µM GDP (Active Rac1/Cdc42 pull-down kit, ThermoFisher, Cat# 16118/19) and incubated for 15 min at 37°C with gentle agitation. Samples of GDP-loaded RhoGTPases were kept on ice for further analyses as controls. Remaining GDP-loaded Rac1 and Cdc42 were then incubated with 500 µM GTPγS (ThermoFisher kit Cat# 16118/19) in the presence or absence of 2 nM of recombinant ERK3 protein (aa 9-327) (Crelux) for 30 min at 37°C at with gentle agitation. All reactions were terminated by addition of 60 mM MgCl_2_ on ice.

To isolate the active (GTP-bound) Rac1 and Cdc42, the specific GST-Pak1-PBD-fusion beads were used. Nucleotide exchange reaction were incubated with the beads for 1 h at 4°C, rotating. Afterwards, beads were washed 3 times with binding/wash buffer: 25mM Tris-HCl, pH 7.2, 150mM NaCl, 5mM MgCl_2_, 1% NP-40 and 5% glycerol. Samples were eluted with 4x sample buffer supplemented with 100 mM DTT. Levels of active Rac1 and Cdc42 were detected using Rac1 and Cdc42 specific antibodies from the active Rac1/Cdc42 pull-down kit (Cat# 16118/19, ThermoFisher). Levels of GST-Pak1-PBD protein were detected using Ponceau S staininbg of the membrane. Efficiency of the GDP-GTP exchange was calculated as ratio of GTPγS+ERK3 samples and GTPγS alone and presented as fold change where GTPγS samples were used as a reference.

Guanine nucleotide exchange (GEF) activity of ERK3 (9-327 aa) was further evaluated using RhoGEF Exchange Assay (Cat# BK100, Cytoskeleton). Assay was performed according to the manufactures’s instructions and included Dbs GEF as a positive control.

#### *In vitro* kinase assay

In order to asses kinase activity of full-lenght (recombinant ERK3 Cat# M31-34G, SignalChem) and kinase domain ERK3 (ERK3 (amino acids (aa) 9-327, Crelux), we performed in vitro kinase assaay using recombinant MK5 (MAPKAPK5 Unactive Cat# M42-14G, SignalChem) and Myelin basic protien (MBP) as substrates (MBP, Dephosphorylated, Cat# 13-110, Merck). For each reaction, 1µg of the selected substrate (MK5 or MBP) was mixed with 0.25 µg of either full-lenght GST-ERK3, GST protein alone (E.coli expressed) or kinase domain ERK3 in kinase buffer (250 mM HEPES pH 7.5, 500 mM NaCl, 25 mM MgCl_2,_ supplemented with protease inhibitor cocktail Set I-Calbiochem 1:100 (Cat# 539131, Merck Millipore) and phosphatase inhibitors: 1 mM sodium orthovanadate (Na_3_VO_4_), 1 mM sodium fluoride (NaF). The MG^2+^/ATP activating solution (Cat# BML-EW9805, Enzo) was added and reations were incubated for 30 min at 30°C on a thermo-mixer with aggitation at 550 rpm. All reactions were stopped by addition of 4x laemmli buffer and boiled at 95°C for 5 min prior Western Blot analyses. For the MBP kinase assay, phospho-Serine and phospho Threonine antibodies were used to assess MBP phosphorylation status. In case of MK5, specific phospho-MK (T182) antibody was used.

#### *In vivo* F-actin/G-actin assay

HMECs and MDA-MB231 cells were seeded in 12-well or 6-well plates for F/G actin assay and parallel Western Blot analyses. When cells reached about 70% confluency, plates were subjected to either Western Blot analyses or G-actin/F-actin *in vivo* assay using cytoskeleton kit (Cat# BK037) according to the manufacturer’s instructions. Briefly, cells were washed with 1xPBS and lysed in appropriate volume of lysis and F-actin stabilization buffer supplemented with 1mM ATP and protease inhibitor mixture for F-actin stabilization (Cytoskeleton). Lysates were centrifuged at 100 000 x g at 37°C for 1 h to separate the G-actin fraction in the supernatants and pelleted F- actin. F-actin pellets were further depolymerized for 1 h on ice using F-actin depolymerizing buffer. Both sets of samples were mixed with 4 x SDS-PAGE sample buffer. Equal volumes of G-actin and F-actin lysates were analysed by SDS-PAGE and immunoblotting with anti-actin rabbit polyclonal antibody (Cat# AANO1, Cytoskeleton). Densitometric analyses of the G-actin and F-actin levels were performed using ImageJ software. Ratios of F-actin in respect to G-actin levels were calculated.

#### *In vitro* actin polymerization assay

Actin polymerization assay was performed using actin polymerization biochem kit (Cat# BK003, Cytoskeleton) according to the provided protocol. Pyrene-labeled rabbit skeletal muscle actin (Cat# AP05-A, Cytoskeleton) (2.3 µM per reaction) was diluted with general actin buffer (5mM Tris-HCl pH 8.0, 0.2 mM CaCl_2_) supplemented with 0.2mM ATP and 1mM DTT and incubated for 1 h on ice to depolymerize actin oligomers. Actin stock was centrifuged for 30 min at 14000 rpm at 4°C. Pyrene-labeled actin was incubated with recombinant ARP2/3 protein complex alone (Cat# RP01P, Cytoskeleton) or along with the human recombinant proteins: WASP-VCA domain protein (Cat# VCG03, Cytoskeleton) or ERK3 protein (M31-34G, SignalChem). ARP2/3 protein complex, WASP-VCA domain protein and full-length ERK3 recombinant protein were used at 10 nM, 400 nM and 4.8 nM final concentration, respectively. Actin alone was used to establish a baseline of polymerization rate. Actin polymerization was induced by addition of 1.5 x actin polymerization buffer (Cat# BSA02) (10x buffer: 20 mM MgCl_2_, 500 mM KCl, 10mM ATP, 50 mM guanidine carbonate in 100 mM Tris-HCl, pH 7.5). Actin polymerization was measured by fluorescence emission at 415 nm over 1 h (30-60 s interval time) at room temperature (RT) with 360 nm excitation wavelength using multimode microplate reader Tecan SPARK^®^. Buffer background signal at each interval was subtracted and relative fluorescence unit (RFU) are depicted.

### Immunofluorescence (IF) and confocal analyzes

Cells cultured on coverslips were fixed in 3.7% formaldehyde (Cat# CP10.1, Roth) for 15 min, followed by washing with PBS pH 7.5 and 3 min permeabilisation using 0.1% Triton X-100 (AppliChem). After washing twice with PBS, cells were blocked with 1% BSA in PBS for 15 min and washed once with PBS. Filamentous actin was labeled with Oregon Green 488 Phalloidin (Cat# O7466, Invitrogen) or alernatively Rhodamine Phalloidin (Cat# R415, Invitrogen) in blocking buffer for 1 h at room temperature in the dark. Nuclei were stained with 10 µg/ml of DNA dye (Hoechst 33342) (Cat# H3570, Invitrogen). For co-staining of endogenous ERK3 and F-actin, cells were incubated in 1:250 dilution of anti-ERK3 antibody (mouse Cat# MAB3196, R&D or rabbit Cat# ab53277, Abcam) in blocking solution for 1 h at RT. For the ARP3 and Cdc42 staining, the anti-ARP3 antibody (Cat# ab151729, Abcam) and anti-Cdc42 antibody (Cat# 610929, BD Transduction) were used in blocking buffer for 1 h at room temperature in the dark. Nuclei were stained with 10 µg/ml of

DNA dye (Hoechst 33342) (Cat# H3570, Invitrogen). Afterwards, cells were washed with PBS and incubated with secondary antibody secondary anti-mouse IgG- Cyanine3 (Cat# A10521, ThermoFisher Scientific) or Ale at 5 µg/ml, DNA dye (Hoechst 33342) and Green Phalloidin in blocking solution for 1 h at RT in the dark. Samples were washed twice with PBS and cells were mounted onto glass slides using Moviol (+DABCO) (Sigma). For exogenous ERK3 detection upon overexpression of the ERK3 WT or serine 189 phosphorylation mutants (S189A/S189D) in shERK3 HMECs, V5-tag-specific antibody was used at 1:100 dilution (Cat # R960-25, Invitrogen) (. Cells were imaged using a Leica DMi8 confocal microscope (63x, oil immersion objective). ERK3-depleted cells were used as a control for the ERK3 staining.

For the colocalization analyses, a mask was created to select excluding nuclei where no colocalization was observed. The ARP3 and ERK3 images were scaled and thresholded in the same way to maximize the dynamic scale and then had the colocalization of the Pearson’s correlation and Spearman’s rank correlation coefficients (PCC and SCC) on a pixel-wise basis with the “Coloc 2” algorithm in Image J. For both the Pearson and Spearman correlation coefficients, a value of +1 is perfectly correlated or colocalized, a value of zero means that the two signals are randomly localizing, and a value of -1 indicates that the two signals are perfectly separated.

### Scanning Electron Microscopy (SEM)

Control and ERK3-depleted HMECs were seeded in 12-well plate. Cells were fixed overnight with 2.5% glutaraldehyde, rinsed with PBS and then again fixed and stained with 2% Osmium Tetroxide. Afterwards, samples were rinsed with distilled water, frozen and then freeze dried with Crist Alpha LSC Plus Freeze Drier. The SEM scans were performed with a Philipps ESEM XL30 scanning electron microscope.

### Filopodia quantification and analyses

To detect, quantify and analyse the number and lengths of the filopodia of the fixed and actin-phalloidin stained control and ERK3 knockdown HMECs, the FiloQuant open access software and routines ^99, 100^ were applied in ImageJ and in some cases were applied in Imaris version 9.3.1 (Imaris, RRID: SCR_007370) for additional verification. Briefly, the raw cell images were linearly adjusted for brightness and contrast for optimal filopodial observation and also had a mask applied to just analyse the filopodial region of interest in each cell and applied to each cell in the same way. Additionally, the following parameters were initially applied to each cell in the same way: Cell Edge Threshold = 20, Number of Iterations = 10, Number of Erode Cycles = 0, Fill Holes on Edges = checked, Filopodia Threshold = 25, Filopodia Minimum Size = 10 pixel. The results of the detected filopodial filaments were the overlayed in white over the actin-phalloidin green fluorescence of the filapodia for visual verification in ImageJ and in some cases in Imaris version 9.3.1.

### Live cell imaging and cell tracking analyses

GFP-actin expressing control and ERK3 knockdown MDA-MB231 cells were cultured in sterile chambers (1×10^4^ cells/well) (µ-slide 8-well, ibidi), coated with 100 µl/well of 5% collagen in DMEM+FBS (Rat tail collagen, Cat# 5056, Advanced BioMatrix) and kept under 5% CO_2,_ 37 °C, and 90% humidity in a OkoLabs environmental incubator (H-301K environmental chamber, Oko Touch, Oko Pump, T-Control and CO2 control, OkoLabs) on the Leica Confocal microscope table.

The images of both GFP fluorescence and transmission of the live cells were acquired with a Leica SP8 confocal microscope using a 10x 0.3 NA objective, with 488 nm excitation (at approximately 150 µW) and emission window of 500-590 nm for GFP detection and with scanning differential interference contrast transmission imaging in a 1500 µm x 1500 µm frame format with 400 lines per second, 0.71 µm/pixel (2048 x 2048 pixel per frame) and with 2 times averaging per line with a frame acquisition of every 30 min per selected position within the chamber. The images were first acquired in multiple regions of each cell type for 20 h.

The image sequences were imported into Imaris version 9.3.1 and detected automatically by fluorescence with both the whole-cell spot and whole-cell surface analysis. As the surface and spot analysis center positions differed in control analysis by less than 1%, the whole cell spot automated analysis was applied to all images in the same way with a 16 µm per cell diameter estimate. The automated tracking also occurred in the same way for all image sequences within Imaris with the autoregressive motion algorithm, with a maximum average distance of 60 µm per step and with zero step gap applied. The speed (or cell speed) was determined by dividing each x, y step by 1800 seconds (or 30 minutes). The acceleration (or cell acceleration) was determined by subtracting a step speed from the previous step speed and dividing by 1800 seconds (or 30 minutes). The displacement length was determined by subtracting the initial x, y position from the final x, y position and then to determine the difference vector length for each track over 20 hours of acquisition. The track length added each absolute x, y vector step for an entire track over 20 hours of acquisition. The track mean speed averaged the speed of the steps for individual tracks.

The graphs were created with GraphPad Prism (RRID:SCR_002798) and cell distribution according to the analyzed parameter was visualized by violin plots. Significance was determined using non-parametric Mann-Whitney test.

### Transwell cell migration assay

Migratory properties of control and ERK3-depleted HMECs and MDA-MB231 cells were assessed using two-chamber transwell system (Cat# 3422, Corning). Cells were seeded in 12-well plates. HMECs were deprived of supplements from the medium 24 h prior the migration. HMECs and MDA-MB231 cells were then trypsinized and resuspended in supplements-free or serum-free medium, respectively. 500 µl of supplements-free/serum-free medium was mixed with 100 ng/ml of EGF and added into the bottom chamber and 1 x 10^5^ cells in 120 µl of supplements-free/serum-free medium were added into each insert. Plates were incubated at 37°C for 24 h. To quantify the migrated cells, cells were removed from the upper surface of the insert using cotton swabs and inserts were washed with PBS. Afterwards, cells that migrated to the lower side were fixed with in 3.7% formaldehyde for 10 min and stained with Hoechst solution in PBS for 20 min at 37°C. Cells were then visualized using Leica DMi8 microscope (5x dry objective), images of three to four regions per each experimental condition were taken. Number of cells was quantified by Fiji/ImageJ software (Fiji, RRID: SCR_002285) using particle analyses for each field of view and averaged for each membrane. Directional migration was presented and the percentage of the respective control (expressed as 100%).

### Phosphorylation site identification on ARP3

Phosphorylation of the ARP2/3 protein complex subunits by full-lenght ERK3 was detected by *in vitro* kinase assay and in-gel digestion of the respective ARP2/3 complex proteins followed by subsequent spectrometry analyses.

#### In-gel digestion

The protein in the gel lanes were digested with 0.1 µg trypsin (Promega) in 20 µl 25 mM ammonium bicarbonate, pH 7.8 at 37°C for 16 h. The tryptic peptides were purified OMIX C18, 10 µl SPE tips (Agilent, Santa Clara, CA, USA), and dried using a Speed Vac concentrator (Savant, Holbrook, NY, USA).

#### Phosphopeptide enrichment

Dried tryptic peptide samples were dissolved in loading buffer (1 M glycolic acid, 6 % trifluoroacetic acid, 5 % glycerol, and 80 % acetonitrile) under continuous shaking. TiO2 beads (Titansphere, TiO2, GL Sciences Inc) were washed in loading buffer three times before transferring them to the dissolved tryptic peptide samples. After 1 h of continuous shaking, the supernatant was collected and transferred to a new tube containing freshly washed TiO2 beads for a second incubation. The TiO2 beads were collected separately and gently washed with 200 μL of loading buffer, 200 μL 80 % acetonitrile/2 % trifluoroacetic acid, 200 mM ammonium glutamate, and 200 μL of 50 % acetonitrile/1 % trifluoroacetic acid, respectively. The TiO2 beads were dried and bound peptides were eluted sequentially in 10 minutes at first with 50 μL of 10 % ammonium hydroxide, pH 11.7, then with 50 μL of 15 % ammonium hydroxide/60 % acetonitrile, and finally with 50 μL of 1 % pyrrolidine. Eluted peptides were acidified by adding 75 μL 50% formic acid and cleaned up using OMIX C18, 10 µl SPE tips (Agilent, Santa Clara, CA, USA).

#### LC-MS analysis

The samples were solved in 10 µl 0.1% formic acid and 5 µl was analyzed by LC-MS using a timsTOF Pro (Bruker Daltonik, Bremen, Germany) which was coupled online to a nanoElute nanoflow liquid chromatography system (Bruker Daltonik, Bremen, Germany) via a CaptiveSpray nanoelectrospray ion source. The peptides were separated on a reversed phase C18 column (25 cm x 75 µm, 1.6 µm, IonOpticks (Fitzroy, VIC, Australia)). Mobile phase A contained water with 0.1% (vol/vol) formic acid, and acetonitrile with 0.1% (vol/vol) formic acid was used as mobile phase B. The peptides were separated by a gradient from 0-35% mobile phase B over 25 min at a flow rate of 300 nl/min at a column temperature of 50°C. MS acquisition was performed in DDA-PASEF mode.

### ERK3-dependent tumor cell motility *in vivo*

All *in vivo* experiments were performed in accordance with the Swiss animal welfare ordinance and approved by the cantonal veterinary office Basel-Stadt. Female NSG mice were maintained in the Department of Biomedicine animal facilities in accordance with Swiss guidelines on animal experimentation. NSG mice are from in-house colonies. Mice were maintained in a sterile controlled environment (a gradual light–dark cycle with light from 7:00 to 17:00, 21–25°C, 45–65% humidity).

For engraftment of MDA-MB231-GFP, 0.5 x 10^6^ cells were suspended in 50 μl Matrigel in PBS (1:1) and injected into the fourth mouse mammary gland of an eight- week-old female. Orthotopic mammary tumors were grown for 4-5 weeks.

### Intravital imaging

Mice were anaesthetized with Attane Isofluran (Provet AG) and anaesthesia was maintained throughout the experiment with a nose cone. Tumors were exposed by skin flap surgery on a Nikon Ti2 A1plus multiphoton microscope and imaged at 880 nm with a Apochromat 25x/1.1NA water immersion objective at a resolution of 1.058 µm per pixel. Cell motility was monitored by time-lapse imaging over 30 min in 2-min cycles, where a 100 mm Z-stack at 5-mm increments was recorded for each field of view starting at the tumor capsule. Three-dimensional time-lapse videos were analysed using ImageJ. Images were registered using demon algorithm in MATLAB to correct for breathing movement (Dirk-Jan Kroon (2022). multimodality non-rigid demon algorithm image registration (https://www.mathworks.com/matlabcentral/fileexchange/21451-multimodality-non-rigid-demon-algorithm-image-registration), MATLAB Central File Exchange). Tumor cell motility was quantified manually. A tumor cell motility event was defined as a protrusion of half a cell length or more over the course of a 30 min video.

### Statistical analyses

All experiments were repeated at least three times and exact replicate number (n) is specified for each figure. GraphPad Prism 9 was used to analyze data. Analyses performed for each data set, including statistical test and post-test were specified for each figure. Where applicable, data are presented as mean ± SEM of at least three independent experiments. Significance levels are displayed in GP (GraphPad) style: *p<0.0332, **p<0.0021, ***p<0.0002, ****p<0.0001.

## Figure supplements

**Figure 1-figure supplement 1.**
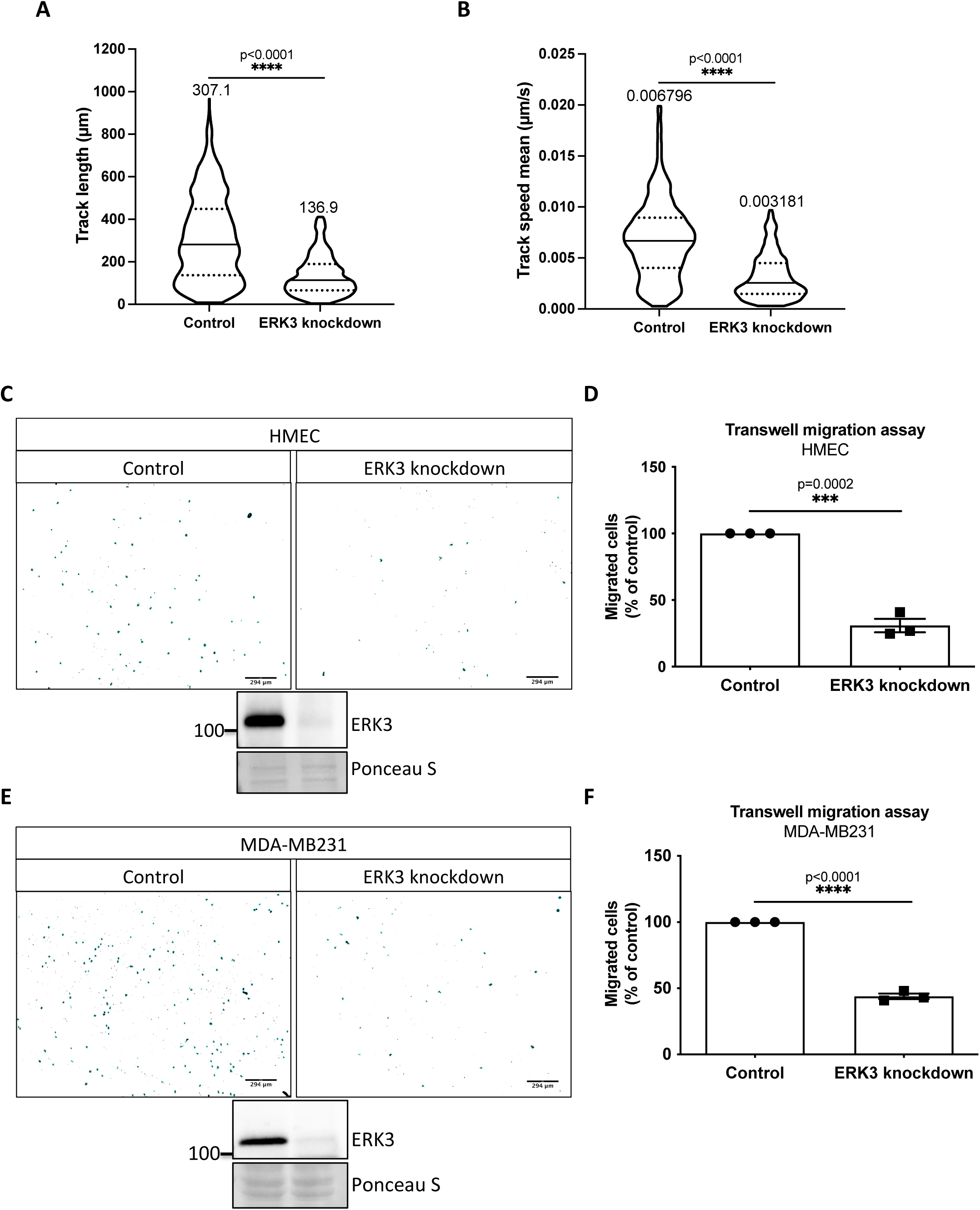
**(A-B)** Tracks analyses depicting **(A)** length (µm) and **(B)** mean speed (µm/s) of 605 random cells (n=605); *p<0.0332, **p<0.0021, ***p<0.0002, **** p<0.0001, Mann-Whitney test. Mean values for each parameter were indicated on top of the violin plots. **(C-F)** Chemotactic responses of control vs. ERK3-depleted **(C-D)** HMECs and **(E-F)** MDA-MB231 cells towards EGF. **(C)** and **(E)** depict representative ImageJ images of the analyzed Hoechst-stained migrated cells and immunoblots of ERK3 knockdown validation. Ponceau S was used as a loading control. **(D)** and **(F)** present comparisons of the chemotactic rate between control and ERK3 knockdown **(D)** HMECs and **(F)** MDA-MB231 cells. Quantification of three (n=3) independent experiments is presented as mean ± SEM percentage of the control; *p<0.0332, **p<0.0021, ***p<0.0002, ****p<0.0001, unpaired t-test.

**Figure 1-figure supplement 2.**
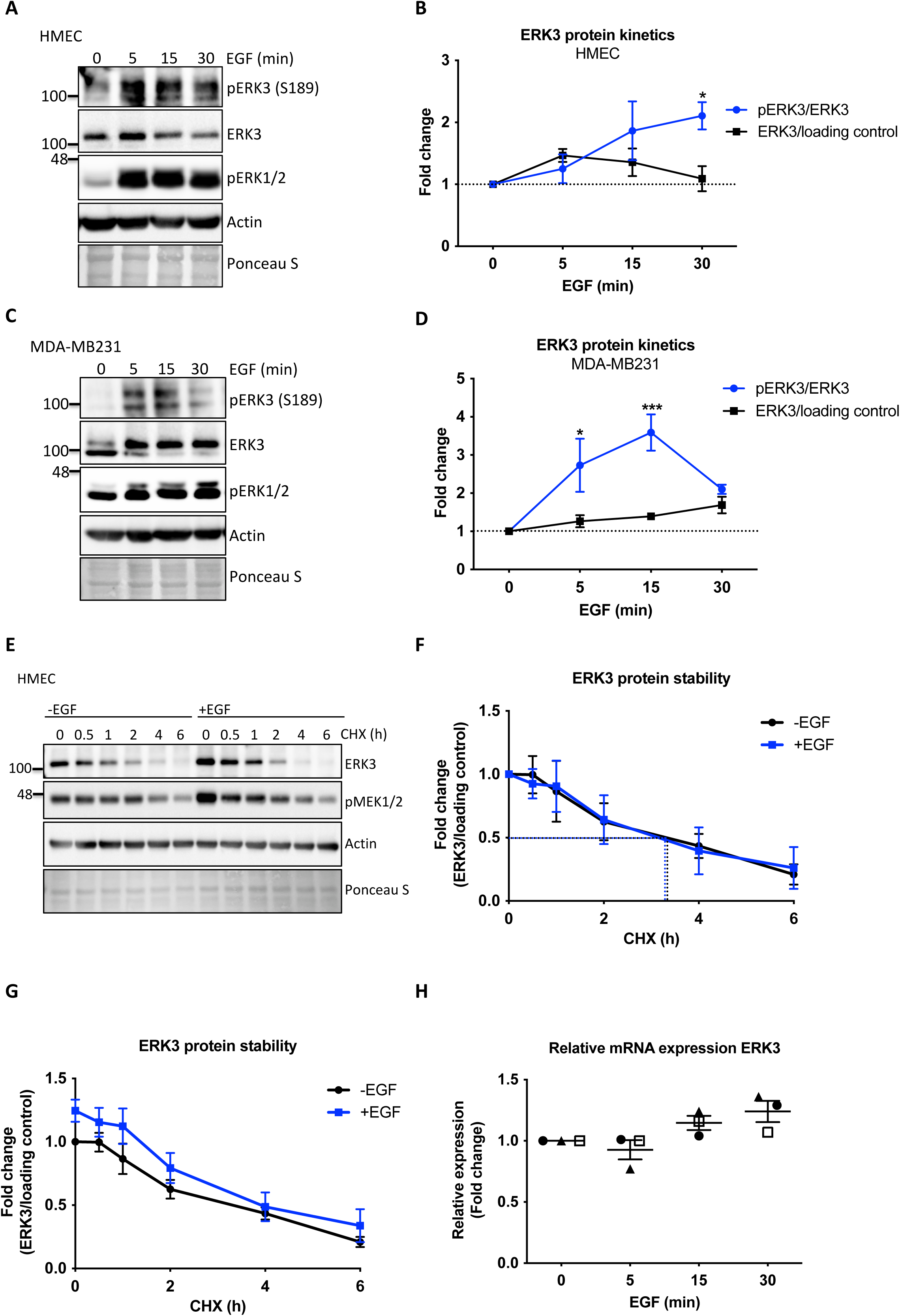
Effect of EGF on chemotactic responses and ERK3 protein kinetics in human mammary epithelial cells. **(A-B)** HMECs and **(C-D)** MDA-MB231 cells were stimulated with EGF at 0, 5, 15 and 30 min. Phosphorylation status of ERK3 at S189 and total protein levels were assessed. Phosphorylation levels of canonical ERK1/2 were used as a validation of EGF treatment. Actin and Ponceau S staining were used as loading controls. **(A, C)** Representative Western Blot analyses. Graphs in **(B)** and **(D)** depict changes in the phosphorylation and expression of ERK3 protein after normalization with total protein levels or internal loading control, respectively. Each time point was normalized to unstimulated cells (0). Mean fold changes ± SEM from three independent experiments (n=3) are presented; *p<0.0332, **p<0.0021, ***p<0.0002, ****p<0.0001, two-way ANOVA, Bonferroni post-test. **(E-H)** Effect of EGF on ERK3 protein stability and expression. **(E-G)** ERK3 protein stability was assessed by CHX chase at 0, 0.5, 1, 2, 4 and 6 h in the control (-EGF) and EGF pre-treated (+EGF) HMECs. **(E)** Representative Western Blot analyses of ERK3 levels. Phosphorylation of MEK1/2 was used as a positive control for EGF treatment. Actin and Ponceau S were used as loading controls. **(F-G)** ERK3 protein stability was quantified and is presented as fold change **(F)** with respect to the untreated cells (-EGF, 0 h) and **(G)** with respect to the respective 0 h time points of the untreated cells (-EGF) and EGF-stimulated (+EGF) cells. Data are presented as mean fold change ± SEM from three independent experiments (n=3). **(H)** Quantitative RT-PCR analysis of *ERK3* expression was assessed in HMECs upon 0, 5, 15 and 30 min of EGF treatment. Each biological replicate was measured in triplicates. Log2 fold change in gene expression is presented as mean ± SEM of three independent experiments (n=3); *p<0.0332, **p<0.0021, ***p<0.0002, ****p<0.0001, one-way ANOVA, Tukey’s post-test.

**Figure 1-figure supplement 3.**
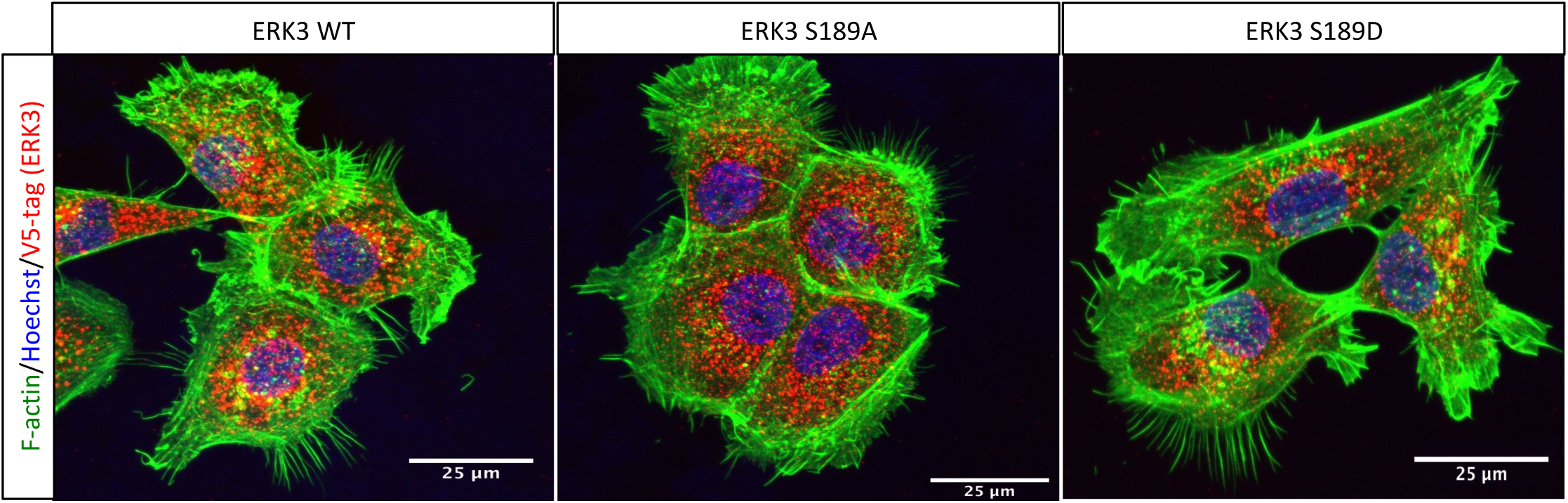
Effect of overexpressed S189 mutants on F-actin. ERK3-depleted HMECs (shERK3, 3’ UTR) were transfected with V5-tagged wild type ERK3 (ERK3 WT) or S189 phosphorylation mutants (ERK3 S189A/ERK3 S189D). Expression of the exogenous ERK3 was determined by staining with V5-tag antibody. Cells were additionally co-stained with phalloidin green and Hoechst to visualize actin cytoskeleton and nuclei, respectively.

**Figure 2-figure supplement 1.**
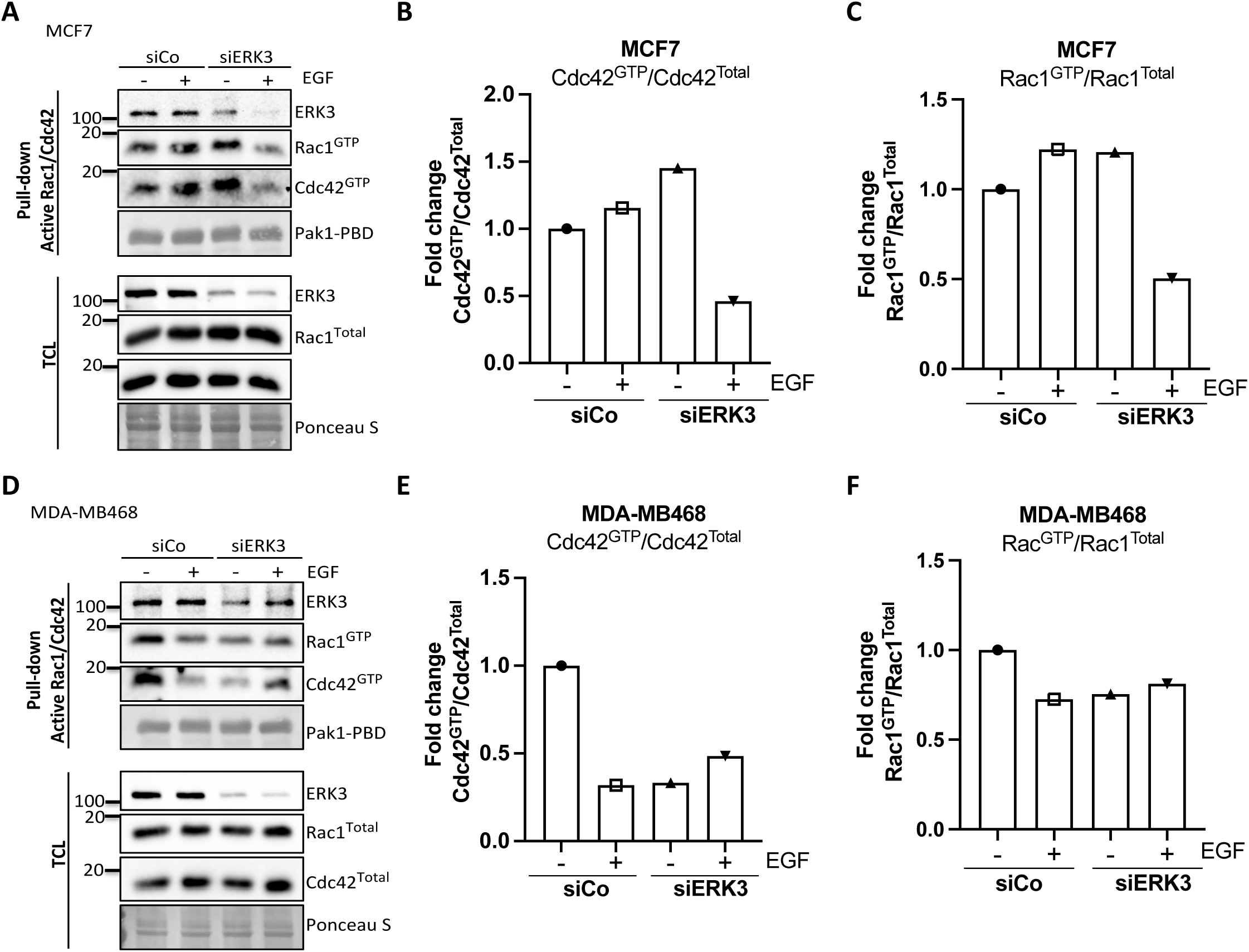
Pull-down of active Rac1 and Cdc42 from control and ERK3-depleted (siERK3 knockdown) breast cancer cells. The siRNA-mediated ERK3 knockdown was introduced in **(A-C)** MCF7 and **(D-F)** MDA-M468 cells. 48 h post-transfection, cells were serum-starved, then treated with EGF (100 ng/ml) for 15 min and subjected to active Rac1/Cdc42 pull-down assay. Levels of active (GTP-bound) Cdc42 and Rac1 as well as co-precipitating ERK3 was assessed in the pulldown. The total protein levels and ERK3 knockdown efficiency were monitored in the total cell lysate (TCL). Relative levels of active Cdc42 **(B and E)** and Rac1 **(C and F)** were calculated with respect to the total protein levels and are presented as mean fold change after normalization with the control (siCo).

**Figure 2-figure supplement 2.**
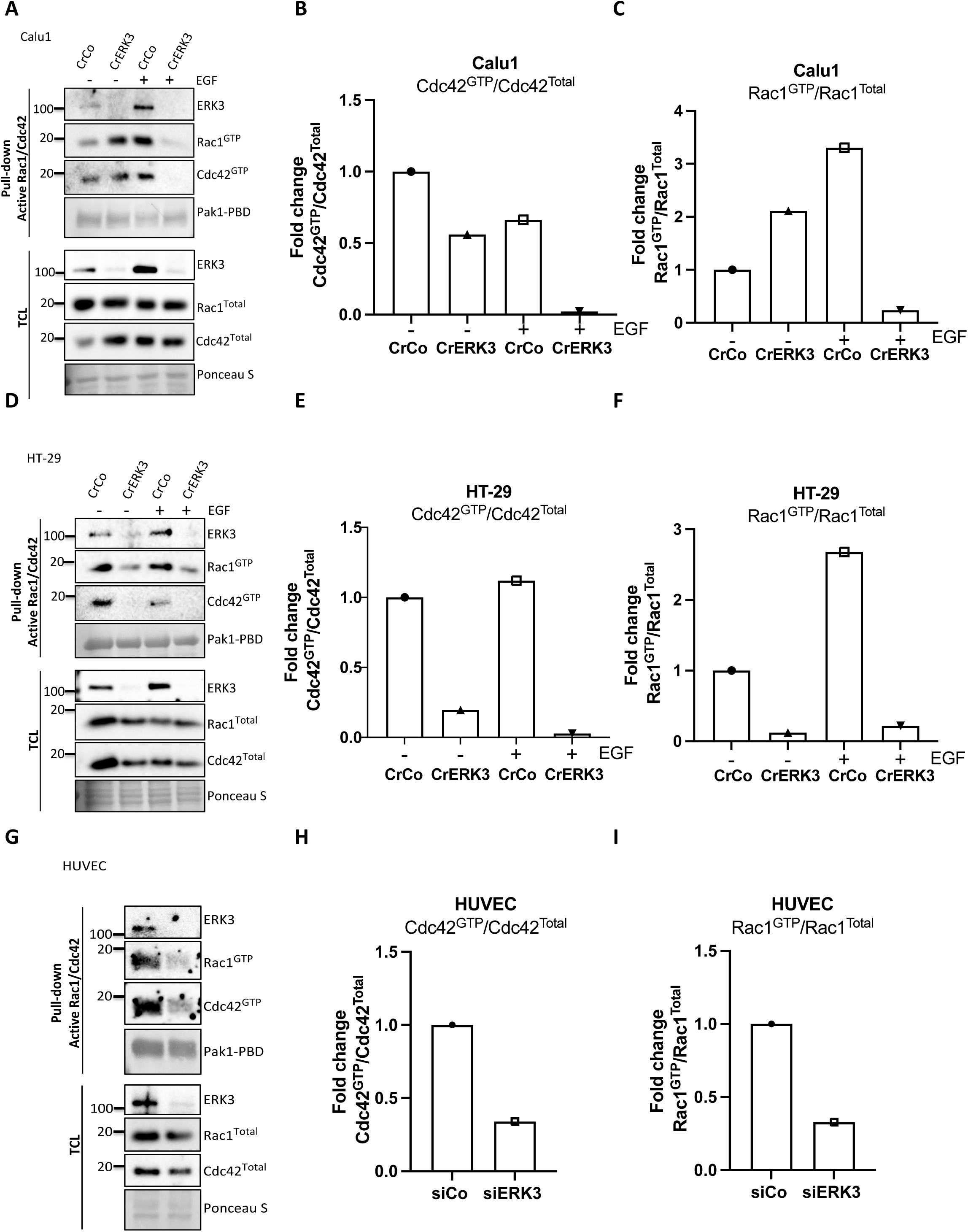
Pull-down of active Rac1 and Cdc42 from CrispR control and CRISPR ERK3 **(A-C)** Calu1 and **(D-F)** HT-29 cells. Serum-starved cells were treated with EGF (100 ng/ml) for 15 min and subjected to active Rac1/Cdc42 pull-down assay. **(G-I)** HUVECs were cultured on 0.2% gelatin-coated plates **and** transfected with siERK3. 48 h post-transfection, cells were subjected to active Rac1/Cdc42 pull-down assay. Levels of active (GTP-bound) Cdc42 and Rac1 as well as co-precipitating ERK3 was assessed in the pulldown. The total protein levels and ERK3 knockdown efficiency were monitored in the total cell lysate (TCL). Relative levels of active Cdc42 **(B, E and H)** and Rac1 **(C, F and I)** were calculated with respect to the total protein levels and are presented as mean fold change after normalization with the control.

**Figure 2-figure supplement 3.**
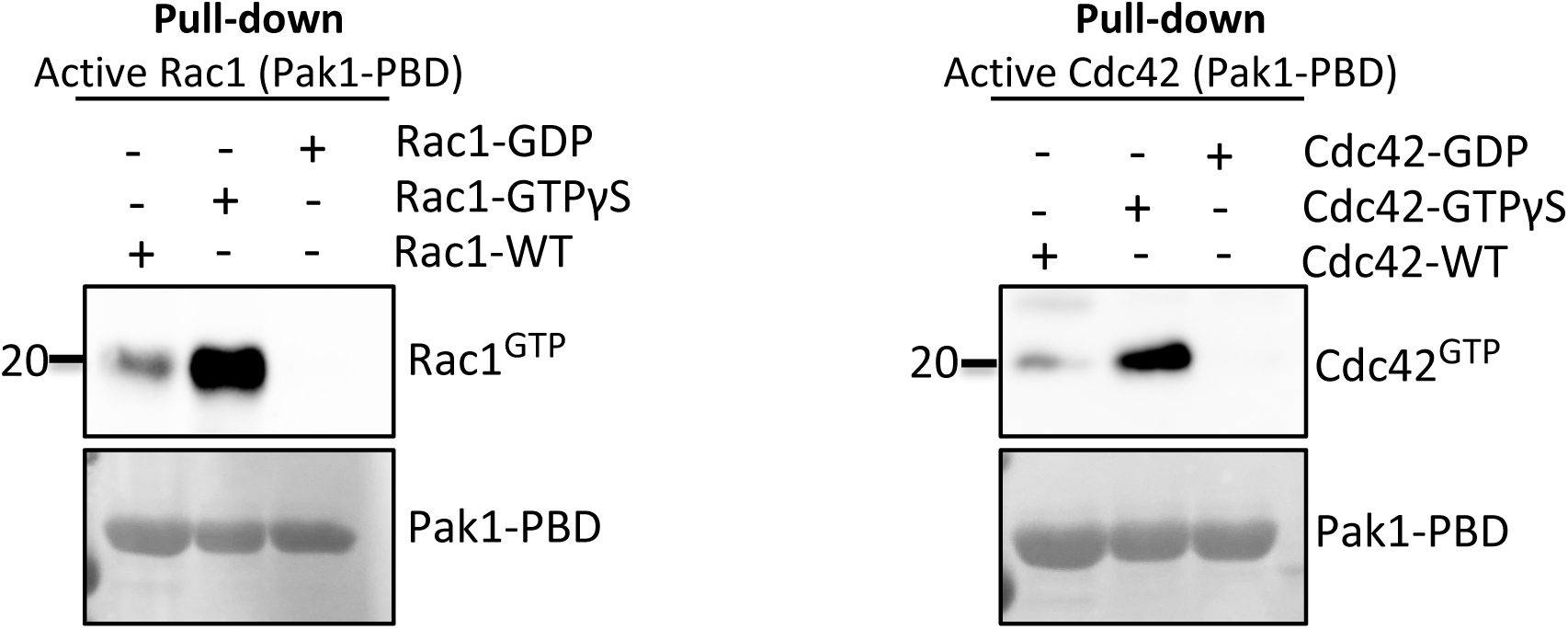
*In vitro* pull-down of purified native and GTP/GTP-loaded Rac1 and Cdc42 proteins using Pak-PBD-fusion beads as described in the methods section. Levels of active (GTP-bound) levels of Rac1 (left panel) and Cdc42 (right panel) were assessed by Western Blot analyses.

**Figure 2-figure supplement 4.**
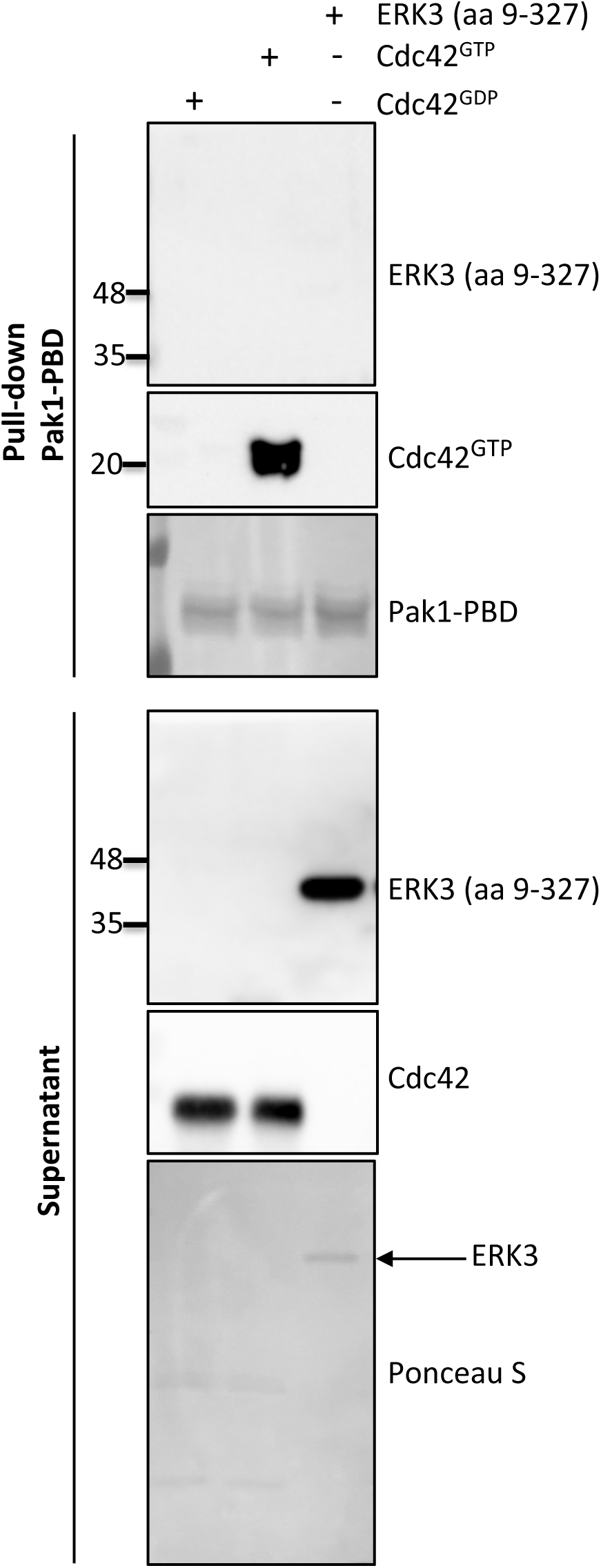
Western Blot analyses of *in vitro* pull-down assay assessing binding affinity of ERK3 (aa 9-327) and Pak1-PBD. Cdc42 was pre-loaded with GTPγ or GDP as described in the methods section and used as positive and negative control for the pull-down, respectively. Pre-loaded Cdc42 or ERK3 (aa 9-327) recombinant proteins were incubated with the Pak1-PBD beads according to the methods section. Levels of active RhoGTPase were detected using Cdc42-specific antibody. Pak1-PBD protein was detected by Ponceau S staining. Supernatant obtained after the pull-down was analyzed to verify the presence of ERK3 and Cdc42 used for the assay. The marked ERK3 protein was also detected with Ponceau S.

**Figure 3-figure supplement 1.**
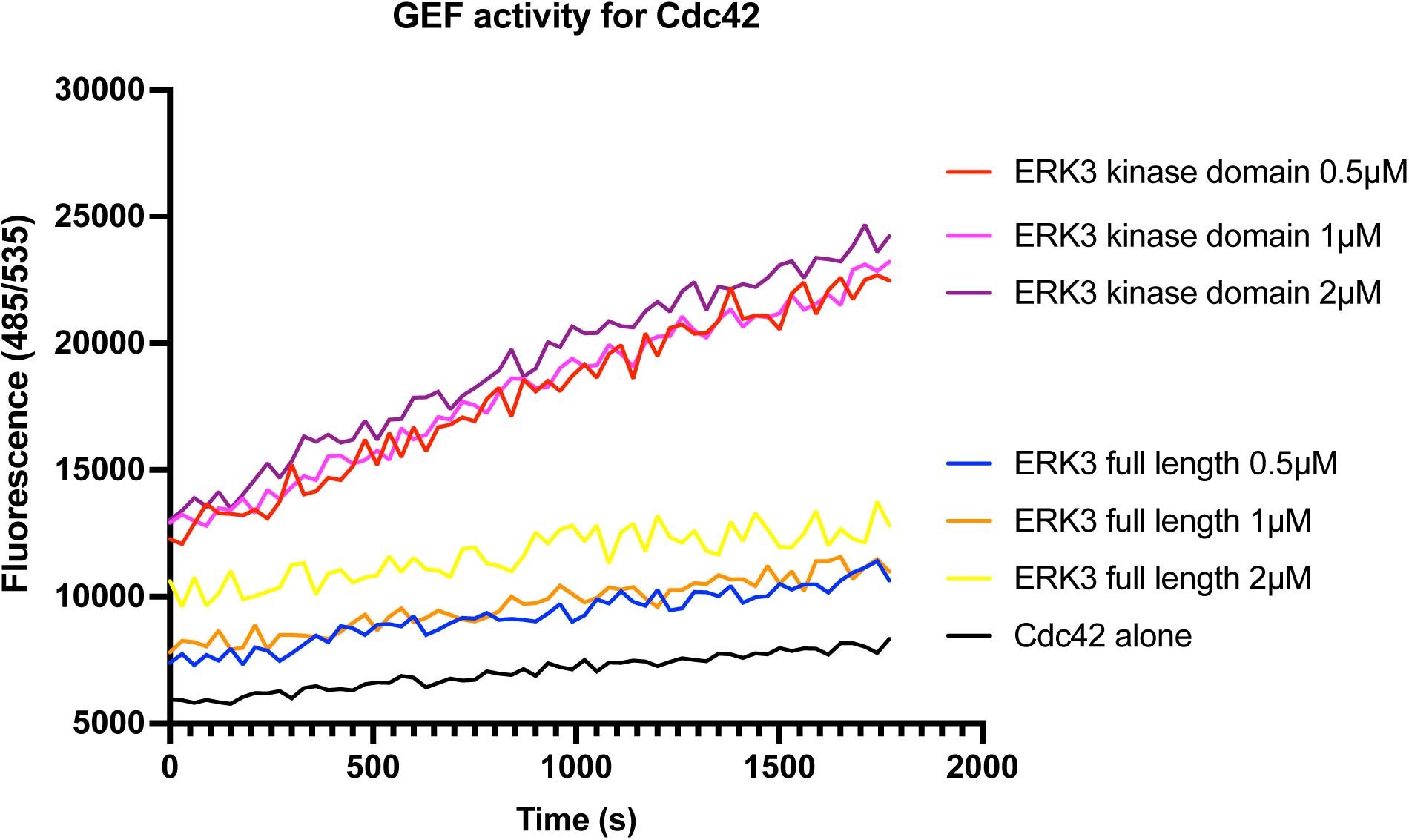
*In vitro* GEF activity of full-length ERK3 vs. ERK3 kinase domain towards Cdc42 was assessed. ERK3 protein were used at 0.5, 1 and 2 µM final concentration. GEF activity was expressed as mean relative fluorescence units (RFU).

**Figure 5-figure supplement 1.**
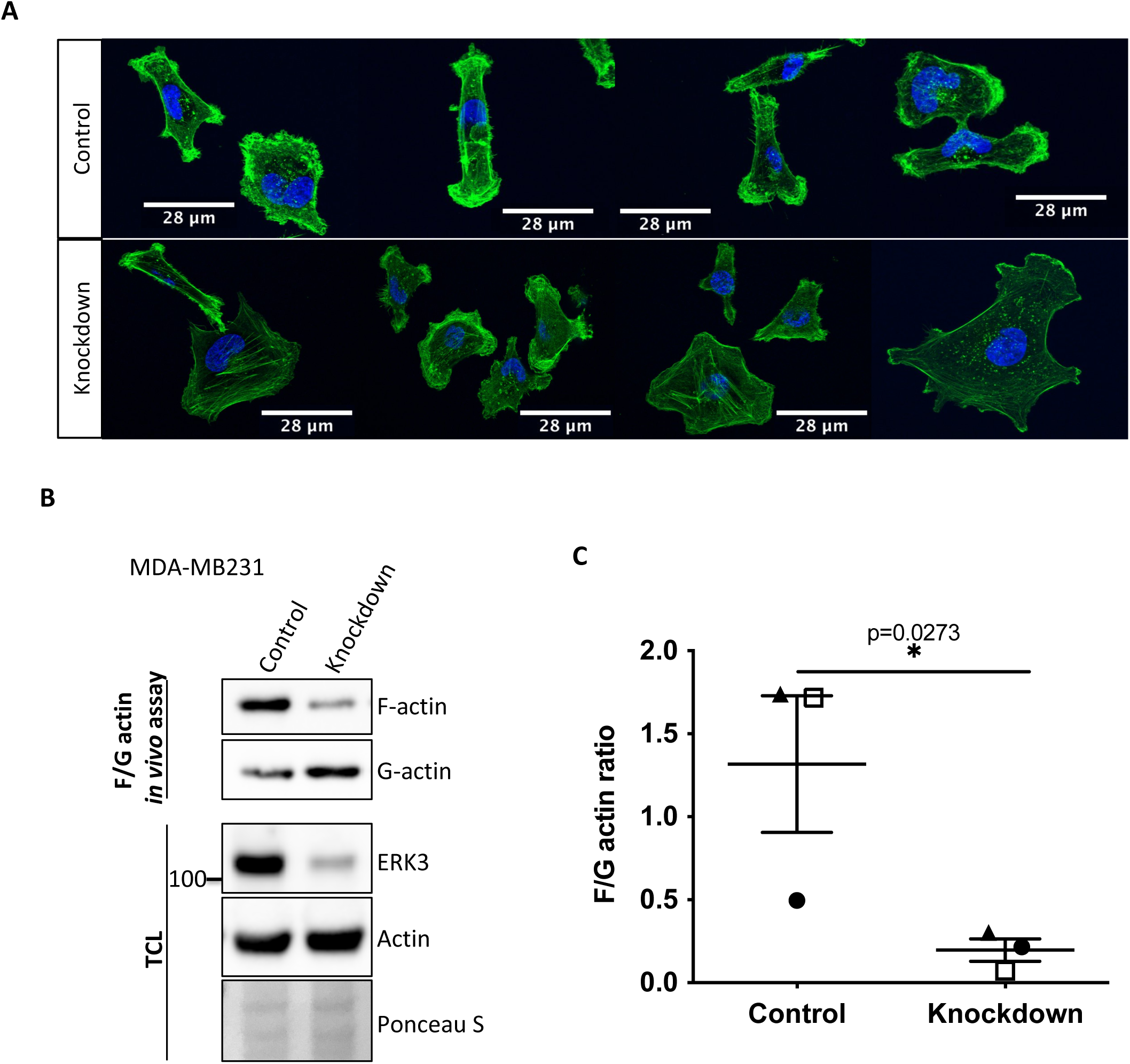
ERK3 regulates F-actin levels in MDA-MB231 cells. F-actin distribution and levels were determined in control (shCo) and ERK3 knockdown (shERK3) MDA-MB231 cells. **(A)** Representative confocal analyzed images of the phalloidin green-stained F-actin merged with Hoechst staining of the nuclei. **(B-C)** Levels of F- and G- actin were quantified using F/G actin *in vivo* assay. **(B)** Representative Western Blot of the F- and G-actin fractions as well as the total cell lysate (TCL) evaluation of the ERK3 knockdown efficiency and total actin levels. Ponceau S was used as loading control.

**Figure 5-figure supplement 2.**
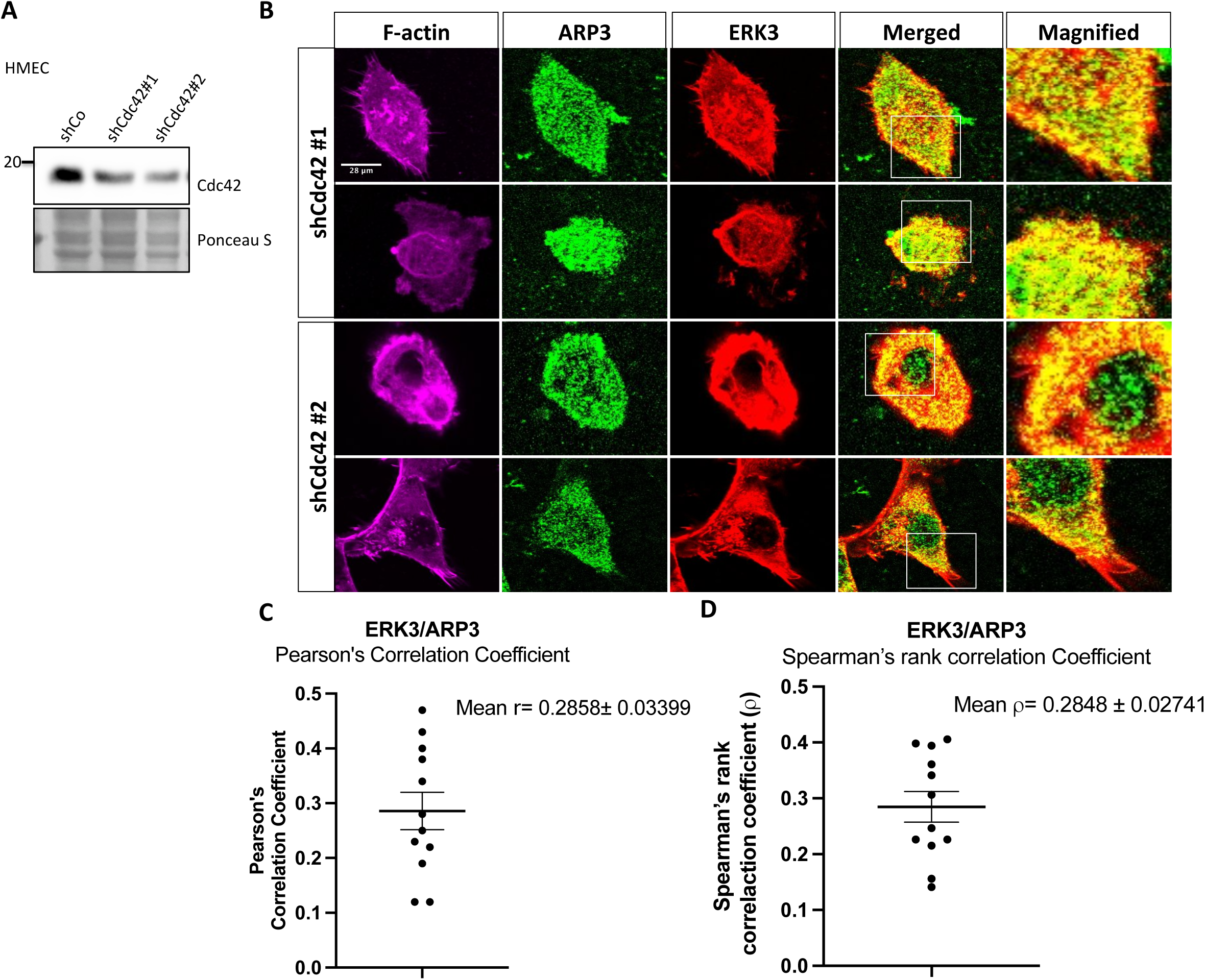
ERK3 and ARP3 (ARP2/3) colocalization in Cdc42- knockdown cells. HMECs were transfected with two shRNAs targeting Cdc42 (shCdc42#1/#2) as described in the methods section. Afterwards, cells were subjected to either **(A)** Western Blot analyses to validate to Cdc42 knockdown or **(B)** IF staining as described in the methods section and confocal imaging to determine localization of ERK3 and ARP3 in the Cdc42 knockdown cells. F-actin was visualized using rhodamine phalloidin to assess cell morphology. **(C-D)** Graphs present **(C)** Pearson’s correlation coefficient and **(D)** Spearman’s rank correlation coefficient values obtained from the co-localization analyses of ERK3 and ARP3 as mean ± SEM from twelve randomly selected cells (n=12).

**Figure 5-figure supplement 3.**
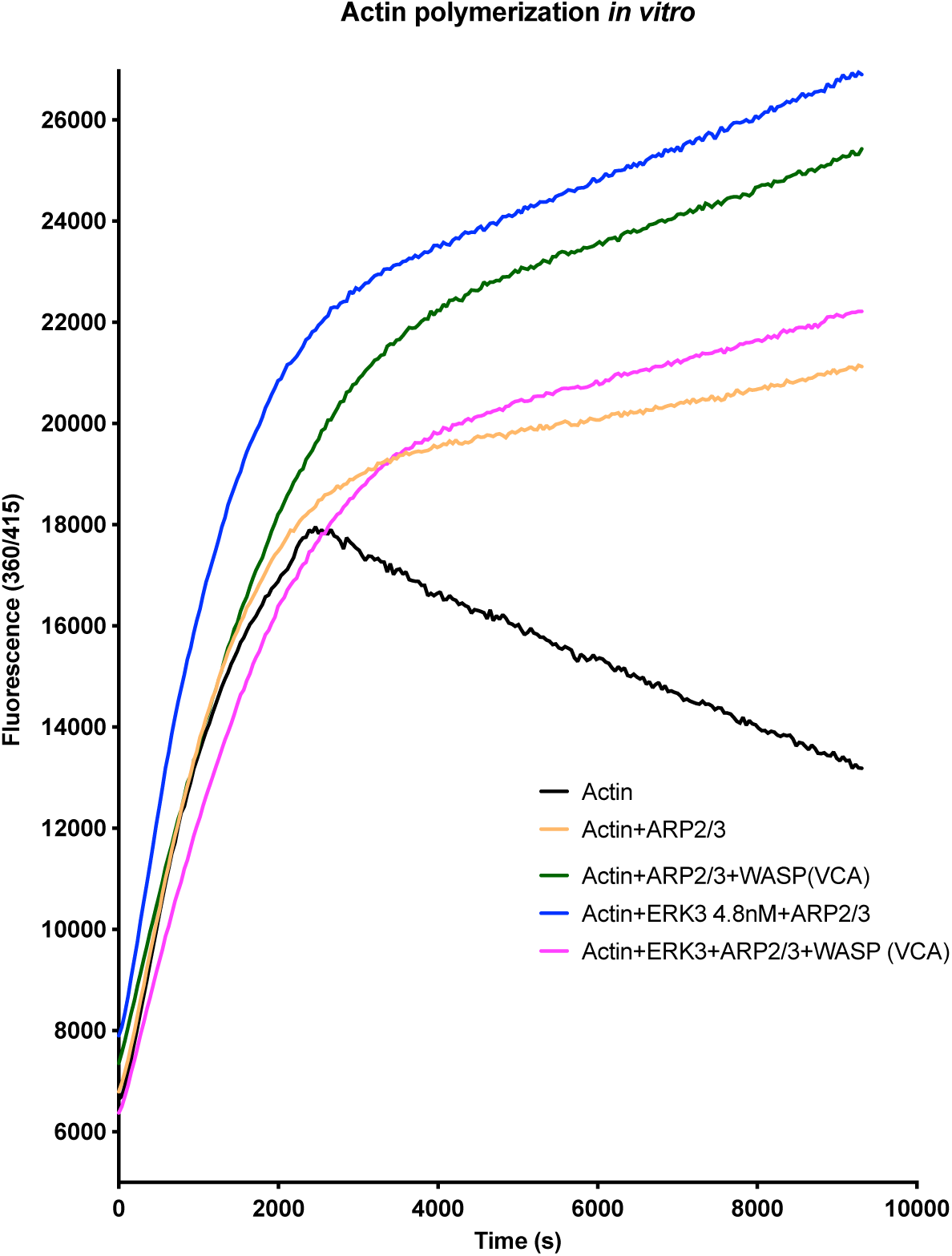
ARP2/3-dependent pyrene actin polymerization *in vitro* was measured in the presence of full-length ERK3 (4.8 nM) or WASP (VCA) domain (400 nM) as well as in the presence of both proteins at the indicated concentrations. Fluorescence at 360/415 was measured over time.

**Figure 6-figure supplement 1.**
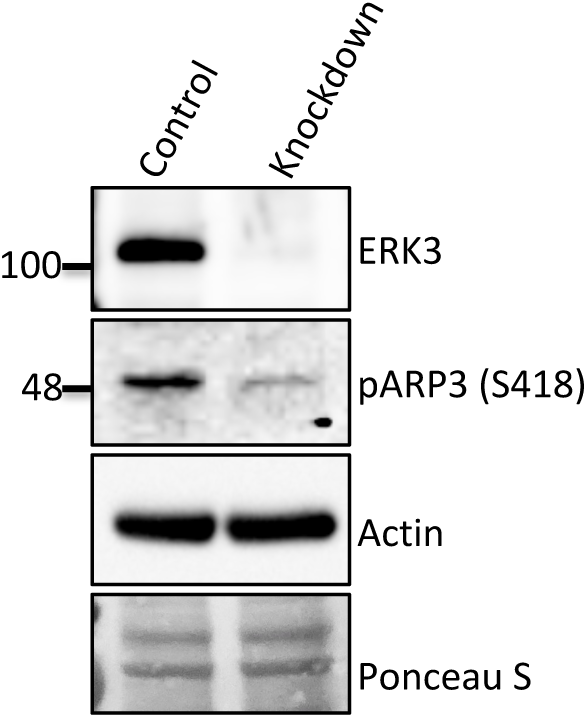
Detection of the S418 phosphorylation of ARP3 in CRISPR ERK3 HMECs presented in Figure 5C-5D).

**Figure 6-figure supplement 2.**
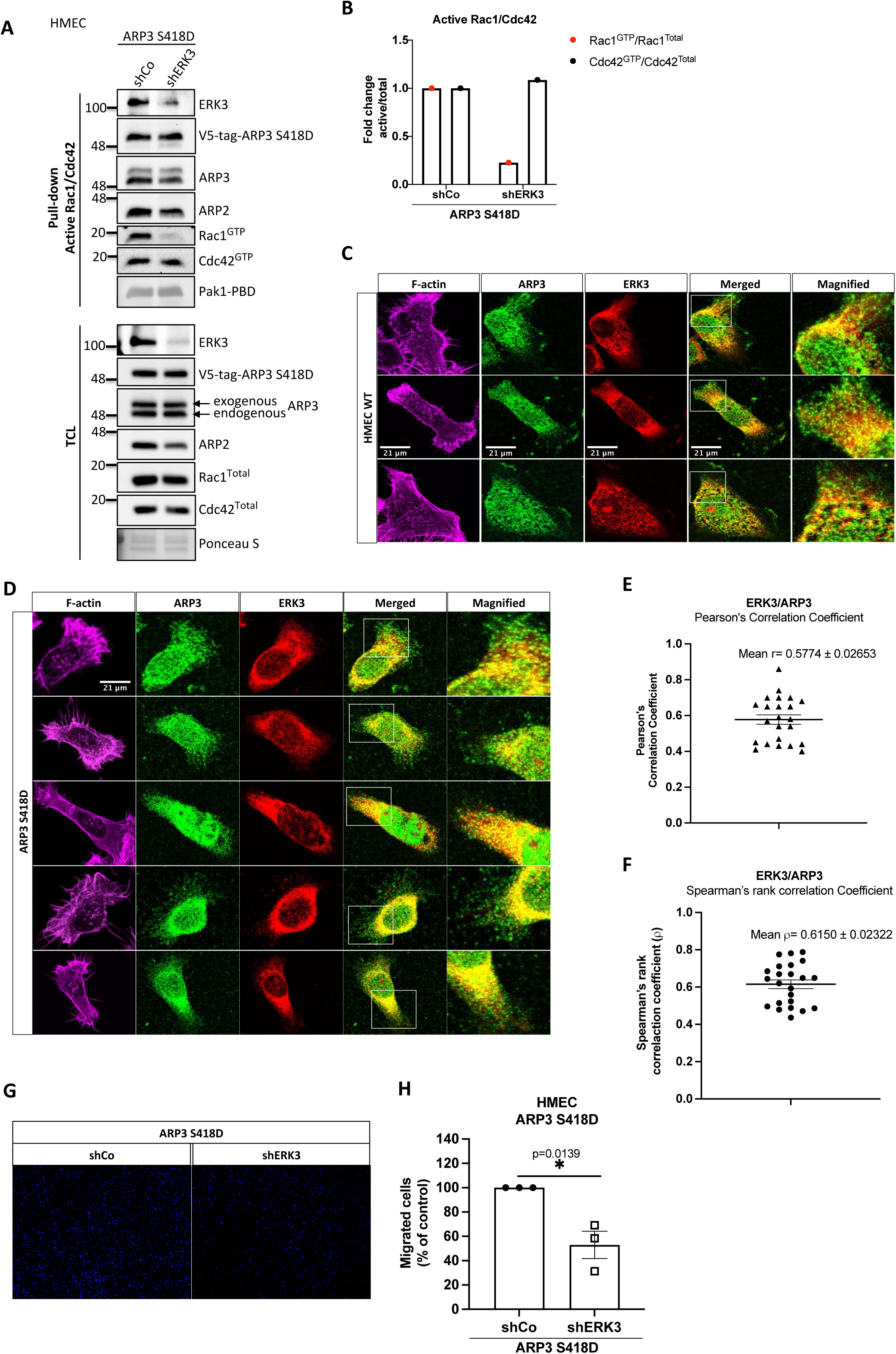
**(A-B)** Active Rac1/Cdc42 pull-down according to manufacturer’s protocol (Cat# 16118/19, ThermoFisher) and methods section. **(A)** Levels of active (GTP-bound) Cdc42 and Rac1 as well as the total protein levels were assessed. Knockdown of ERK3 was validated by ERK3 antibody, exogenous ARP3 (S418D) was detected using a V5-tag antibody and total expression of ARP3 in the cells was determined using an ARP3 antibody. Detection of ARP2 was used as an additional control for the detection of the ARP2/3 complex in both active Rac1/Cdc42 pull-down and TCL. **(B)** Relative levels of active Cdc42 and Rac1 were calculated with respect to the total protein levels and are presented as mean fold change after normalization with the control (shCo). **(C-D)** IF staining of ERK3 (secondary antibody: anti-mouse Alexa Fluor 647 (Cat# A21235, ThermoFisher Scientific)) and ARP3 (secondary antibody: Alexa Fluor 488 (Cat# A11008, ThermoFisher Scientific) in **(C)** WT and **(D)** ARP3 S418D-overexpressing HMECs. F-actin was visualized using rhodamine phalloidin to assess cell morphology. Scale bar: 21 µm. **(E-F)** Graphs present **(E)** Pearson’s correlation coefficient and **(F)** Spearman’s rank correlation coefficient values obtained from the co-localization analyses of ERK3 and ARP3 as mean ± SEM from twenty-three randomly selected cells (n=23). **(G-H)** Effect of the ERK3 knockdown on the directional migration of ARP3 S418D- overexpressing HMECs was assessed and quantified using transwell as described in the methods section. Cells were seeded in the inserts in medium without supplements for 1h prior the beginning of the assay. Complete medium was used as a chemoattractant in the lower chamber. **(G)** Representative images of the analyzed inserts. **H)** Percentage of the migrated ARP3 S418D shERK3 cells as compared to the control (shCo) is presented as mean ± SEM from three (n=3) independent experiments; *p<0.0332, **p<0.0021, ***p<0.0002, ****p<0.0001, t-test.

**Figure 7-figure supplement 1.**
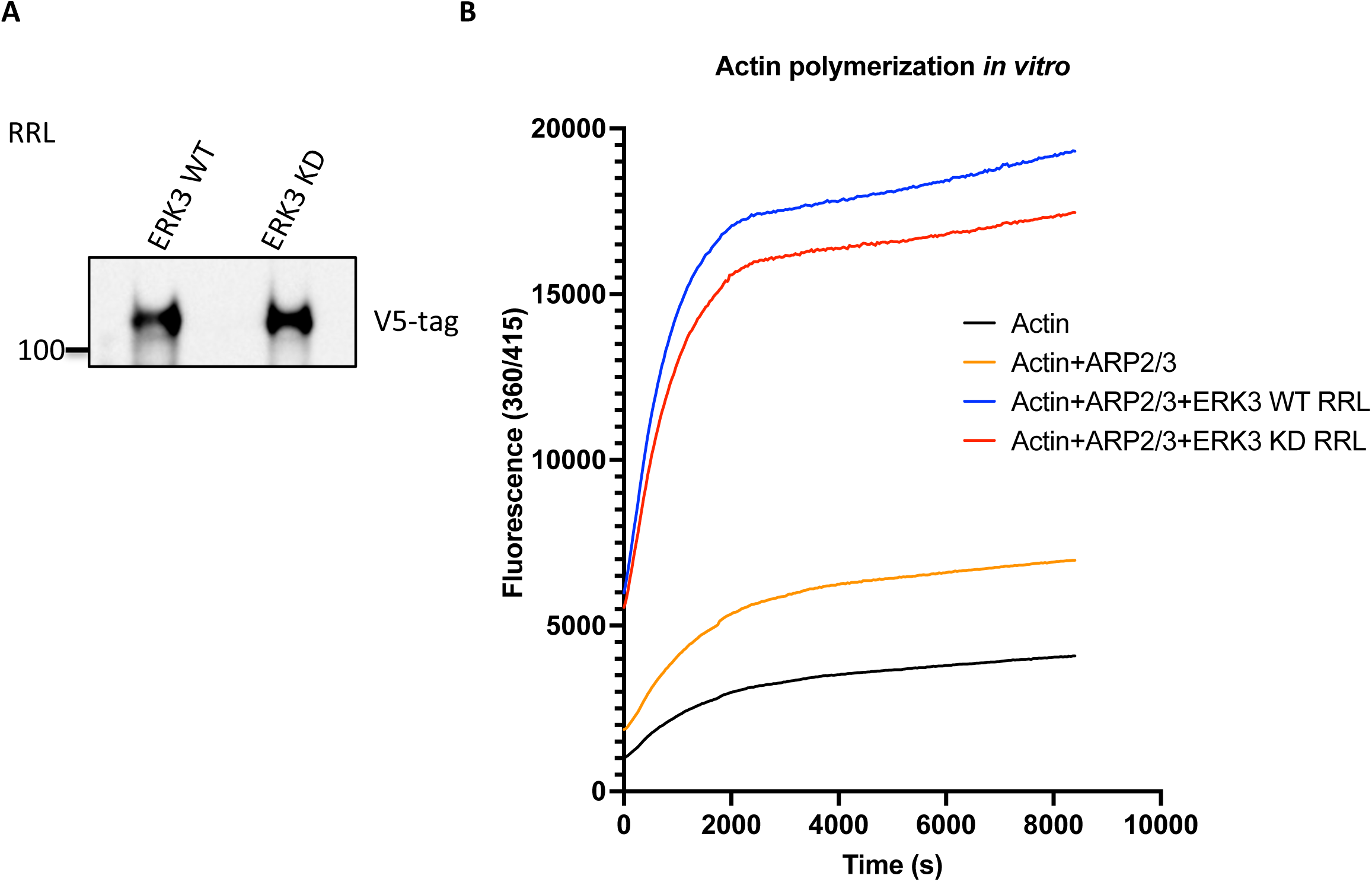
**(A) Western Blot analysis of the RRL-expressed ERK3 proteins. (B)** ARP2/3-dependent pyrene actin polymerization *in vitro* was assessed in the presence of wild type (WT) or kinase-dead (KD) ERK3 proteins expressed in rabbit reticulocyte lysates (RRL). Actin alone baseline was measured in the presence of the same volume of the control RRL as for the expressed proteins. Fluorescence at 360/415 was measured over time and results are shown as mean fluorescence from three experiments.

**Figure 7-figure supplement 2.**
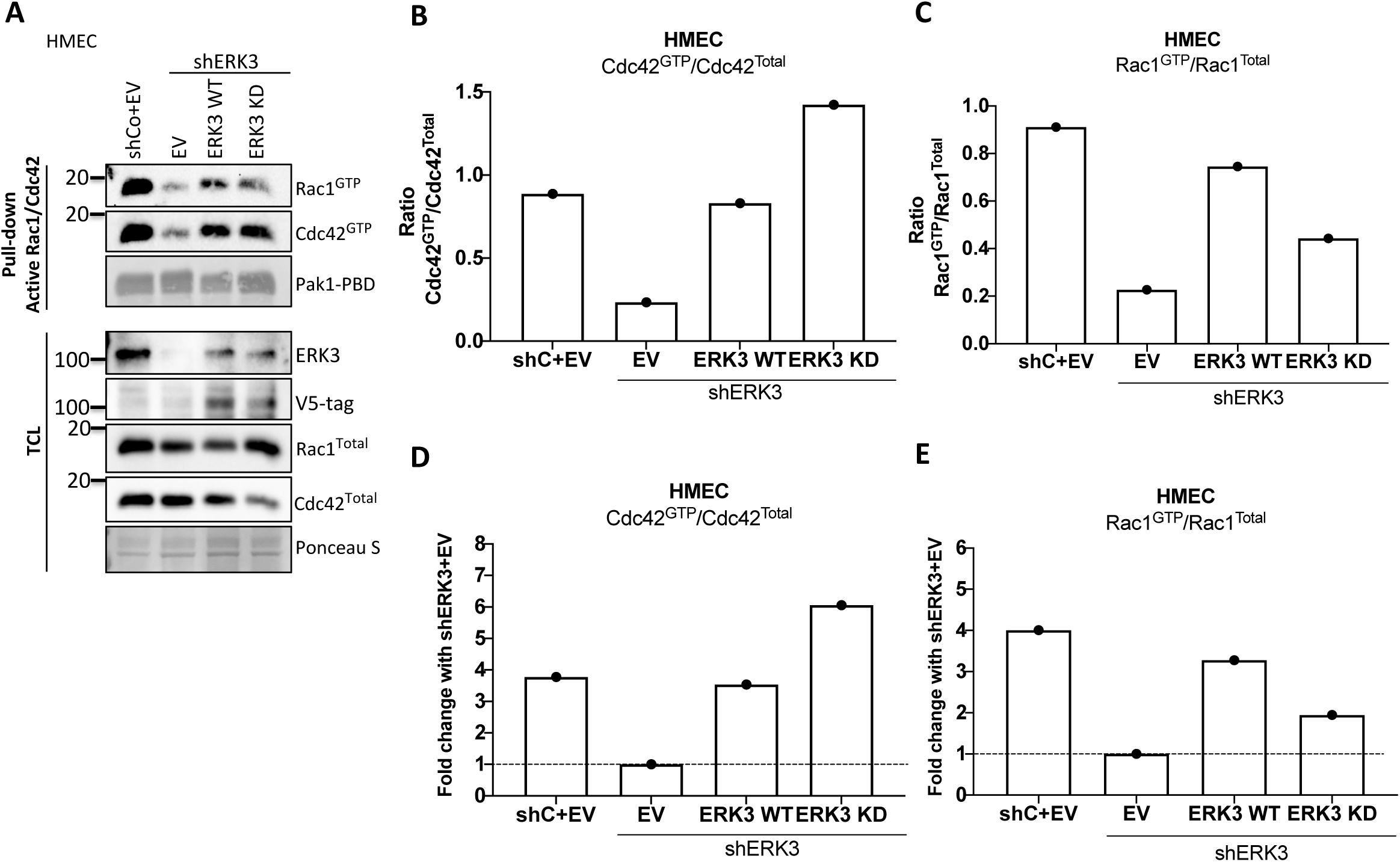
Active Rac1 and Cdc42 pull-down assays were performed from control (shCo) and ERK3-depleted (shERK3, 3’ UTR) HMEC reconstituted with either wild type (WT) or kinase-dead (KD) ERK3 as described in the methods section. Pak1-PBD was used to capture the active form of Cdc42 and Rac1 in the respective cell lysates. Levels of active (GTP-bound) Cdc42 and Rac1 as well as the total protein expression were assessed, ERK3 knockdown efficiency and overexpression was verified in the total cell lysate (TCL) using ERK3 antibody and V5-tag antibody to detect the exogenous WT and KD version of ERK3. **(A)** Western Blot analyses are presented. **(B-C)** Relative levels of active Cdc42 and Rac1 were calculated with respect to the total protein levels and are presented as ratio of GTP/Total RhoGTPases. **(D-E)** Additionally, the ratio of GTP/total Rac1 and Cdc42 was normalized to the control cells (shERK3+EV) and are presented as fold change in activation.

## Supplementary movies

**Movie 1-4.** Movies representing motility of the control (Movie 1 and 2) and ERK3- knockdown (Movie 3 and 4) MDA-MB231 cells.

**Movie 5 and 6.** Representative videos following the motility of the control (shCo) and ERK3 knockdown (shERK3) GFP-expressing MDA-MB231 cells.

## Source data

**Figure 1C-1E-source data.** Prism and Excel file for Figures 1C-1E.

**Figure 1F-source data.** Excel file for Figure 1F.

**Figure 1G-source data.** Prism and Excel file for Figure 1G.

**Figure 1I-source data.** Full membrane scans for Western Blot images for Figure 1I.

**Figure 1J-source data.** Prism and Excel file for Figure 1J.

**Figure1-figure supplement 1A and 1B-source data.** Prisma file and original movie files for Figure 1-figure supplement 1A and 1B.

**Figure 1-figure supplement 1C and 1E-source data.** Full membrane scans for Western Blot images for Figure 1-figure supplement 1C and 1E.

**Figure 1-figure supplement 1D-source data.** Prism and Excel file for Figure 1-figure supplement 1D.

**Figure 1-figure supplement 1F-source data.** Prism and Excel file for Figure 1-figure supplement 1F.

**Figure 1-figure supplement 2A,2C and 2E-source data.** Full membrane scans for Western Blot images for Figure 1-figure supplement 2A, 2C and 2E.

**Figure 1-figure supplement 2B-source data.** Prism and Excel file for Figure 1-figure supplement 2B.

**Figure 1-figure supplement 2D-source data.** Prism and Excel file for Figure 1-figure supplement 2D.

**Figure 1-figure supplement 2F and 2G-source data.** Prism and Excel file for Figure 1-figure supplement 2F and 2G.

**Figure 1-figure supplement 2H-source data.** Prism and Excel file for Figure 1-figure supplement 2H.

**Figure 2B, 2C, 2H, 2I and 2L-source data.** Full membrane scans for Western Blot images for Figure 2B, 2C, 2H, 2I and 2L.

**Figure 2D-source data.** Prism and Excel file for Figure 2D.

**Figure 2-figure supplement 1A, 1D-source data.** Full membrane scans for Western Blot images for Figure 2-figure supplement 1A, 1D.

**Figure 2-figure supplement 1B and 1C-source data.** Prism and Excel file for Figure 1-figure supplement 1B and 1C.

**Figure 2-figure supplement 1E and 1F-source data.** Prism and Excel file for Figure 1-figure supplement 1E and 1F.

**Figure 2-figure supplement 2A, 2D and 2G-source data.** Full membrane scans for Western Blot images for Figure 2-figure supplement 2A, 2D and 2G.

**Figure 2-figure supplement 2B and 2C-source data.** Prism and Excel file for Figure 2-figure supplement 2A and 2C.

**Figure 2-figure supplement 2E and 2F-source data.** Prism and Excel file for Figure 2-figure supplement 2E and 2F.

**Figure 2-figure supplement 2H and 2I-source data.** Prism and Excel file for Figure 2-figure supplement 2H and 2I.

**Figure 2-figure supplement 3-source data.** Full membrane scans for Western Blot images for Figure 2-figure supplement 3.

**Figure 2-figure supplement 4-source data.** Full membrane scans for Western Blot images for Figure 2-figure supplement 4.

**Figure 3B and 3D-source data.** Full membrane scans for Western Blot images for Figure 3B and 3D.

**Figure 3C -source data.** Prism and Excel file for Figure 3C.

**Figure 3E -source data.** Prism and Excel file for Figure 3E.

**Figure 3-figure supplement 1-source data.** Prism and Excel file for Figure 3-figure supplement 1.

**Figure 4A, 4B and 4C-source data.** Full membrane scans for Western Blot images for Figure 4A, 4B and 4C.

**Figure 4F and 4G-source data.** Prism and Excel file for Figure 4F and 4G.

**Figure 5B, 5C and 5D-source data.** Full membrane scans for Western Blot images for Figure 5B, 5C and 4D.

**Figure 5E-source data.** Prism and Excel file for Figure 5E..

**Figure 5F-source data.** Prism and Excel file for Figure 5F.

**Figure 5-figure supplement 1B-source data.** Full membrane scans for Western Blot images for Figure 5-figure supplement 1B.

**Figure 5-figure supplement 1C-source data.** Prism and Excel file for Figure 5-figure supplement 1C.

**Figure 5-figure supplement 2A-source data.** Full membrane scans for Western Blot images for Figure 5-figure supplement 2A.

**Figure 5-figure supplement 2C and 2D-source data.** Prism and Excel file for Figure 5-figure supplement 2C and 2D.

**Figure 5-figure supplement 3-source data.** Prism and Excel file for Figure 5-figure supplement 3.

**Figure 6A, 6C, 6E, 6G, 6H and 6K-source data.** Full membrane scans for Western Blot images for Figure 6A, 6C and 6E, 6G, 6H and 6K.

**Figure 6B-source data.** Prism and Excel file for Figure 6B.

**Figure 6D-source data.** Prism and Excel file for Figure 6D.

**Figure 6I-source data.** Prism and Excel file for Figure 6I.

**Figure 6L-source data.** Prism and Excel file for Figure 6L.

**Figure 6-figure supplement 1-source data.** Full membrane scans for Western Blot images for Figure 6-figure supplement 1.

**Figure 6-figure supplement 2A-source data.** Full membrane scans for Western Blot images for Figure 6-figure supplement 2A.

**Figure 6-figure supplement 2B-source data.** Prism and Excel file for Figure 6-figure supplement 2B.

**Figure 6-figure supplement 2E and 2F-source data.** Prism and Excel file for Figure 6-figure supplement 2E and 2F.

**Figure 6-figure supplement 2H-source data.** Prism and Excel file for Figure 6-figure supplement 2H.

**Figure 7A and 7D -source data.** Full membrane scans for Western Blot images for Figure 7A and 7D.

**Figure 7C-source data.** Prism and Excel file for Figure 7-figure supplement 2H.

**Figure 7-figure supplement 1A-source data.** Full membrane scans for Western Blot images for Figure 7-figure supplement 1A.

**Figure 7-figure supplement 1B-source data.** Prism and Excel file for Figure 7-figure supplement 1B.

**Figure 7-figure supplement 2A-source data.** Full membrane scans for Western Blot images for Figure 7-figure supplement 2A.

**Figure 7-figure supplement 2B-2D-source data.** Prism and Excel file for Figure 7-figure supplement 2B-2D.

## Acknowledgements

The authors thank Kerstin Bahr, Institute of Functional and Clinical Anatomy, University Medical Center of the Johannes Gutenberg University, Mainz for electron microscopy imaging. We would like to thank Stefanie Wenzel and Lisa Winkler for the technical assistance. This work is supported from the grants from Else Kroener Fresenius Stiftung; MERCK (project ID-ERK-KR). We would like to thank the Translational oncology team of MERCK for their valuable inputs. We would like to thank Dr. Raphael Thierry and Dr. Ewelina Bartoszek for their assistance with MATLAB script used for the analyses of the tumor cell motility analyses. We would like to thank Dr. Ulrike Theisen and Prof. Reinhard W. Köster from the department of Cellular and Molecular Neurobiology, of Technische Universität Braunschweig, Braunschweig, Germany for critical reading of the mansucript and valuable input. We thank Dr. Daniela Hoeller for the critical reading and editing of this manuscript.

## Authors contributions

K.B-J: Conceptualization, Methodology, Validation, Formal Analysis, Investigation, Writing-Original Draft, Writing-Review & Editing, Visualization; G.H: Software and Data Curation for the motility experiment; MMC and MB :Methodology, validation, Analysis, Data curation. B.T: Data Curation, Formal Analysis of MassSpec data and Writing-Review & Editing; K.R Conceptualization, Resources, Supervision, Project management Funding Acquisition and Writing-Review & Editing.

